# Myopathy mutations in DNAJB6 slow conformer specific substrate processing that is corrected by NEF modulation

**DOI:** 10.1101/2021.12.22.473881

**Authors:** Ankan K. Bhadra, Michael J. Rau, Jil A. Daw, James A.J. Fitzpatrick, Conrad C. Weihl, Heather L. True

**Affiliations:** Department of Cell Biology and Physiology, Washington University School of Medicine, 660 South Euclid Avenue, Campus Box 8228, St. Louis, MO, 63110, USA. Electronic address; Washington University Center for Cellular Imaging (WUCCI), Washington University School of Medicine, St. Louis, United States; Department of Neuroscience, Washington University School of Medicine, St. Louis, United States; Department of Neurology, Hope Center for Neurological Diseases, Washington University School of Medicine, St. Louis, MO, USA

## Abstract

Molecular chaperones, or heat shock proteins (HSPs), protect against the toxic misfolding and aggregation of proteins. As such, mutations or deficiencies within the chaperone network can lead to disease. In fact, dominant mutations in DNAJB6 (Hsp40/Sis1), an Hsp70 co-chaperone, leads to a protein aggregate myopathy termed Limb-Girdle Muscular Dystrophy Type D1 (LGMDD1). DNAJB6 client proteins and co-chaperone interactions in skeletal muscle are not known. Here, we used the yeast prion model client in conjunction with *in vitro* chaperone activity assays to gain mechanistic insights, and found that LGMDD1 mutants affect Hsp40 functions. Strikingly, the mutants changed the structure of client protein aggregates, as determined by altered distribution of prion strains. They also impair the Hsp70 ATPase cycle, dimerization, and substrate processing and consequently poison the function of wild-type protein. These results define the mechanisms by which LGMDD1 mutations alter chaperone activity and provide avenues for therapeutic intervention.

## Introduction

Limb-Girdle Muscular Dystrophies (LGMDs) are a genetically heterogeneous family of muscle disorders that are either as autosomal dominant or recessive^1^. Although most recessive LGMDs are characterized by a loss-of-function, the mechanistic nature of dominantly inherited LGMDs is unclear. These late onset degenerative myopathies are unified by similar myopathologies that include myofibrillar disorganization, impaired protein degradation, and the accumulation of protein inclusions that contain structural muscle proteins such as desmin and α-actinin and RNA binding proteins such as TDP-43^2–4^. This toxic misfolding and aggregation of proteins is protected by the activity of molecular chaperones, or heat shock proteins (HSPs). As such, mutations in these chaperones can lead to diseases termed “chaperonopathies”. One such example is Limb-Girdle Muscular Dystrophy Type D1 (LGMDD1), caused by mutations in DNAJB6 (Hsp40), an Hsp70 co-chaperone^4^. The originally identified LGMDD1 disease mutations are present within the 12 amino acid stretch of the glycine/phenylanine (G/F) rich domain^3–7^. DNAJB6 is expressed ubiquitously and participates in protein folding and disaggregation^8–11^; however, its role in skeletal muscle protein homeostasis is unknown. In addition, DNAJB6 client proteins and DNAJB6 chaperone interactions in skeletal muscle are not known. Fortunately, DNAJB clients are well-characterized in yeast and thereby afford a model system to study the effect of disease-causing mutants. Moreover, the understanding of DNAJB function within the yeast chaperone network is more complete than in skeletal muscle. Previously, we utilized a transdisciplinary approach to ascertain the functionality of LGMDD1-associated mutants in model systems^12^. Here, we generated homologous DNAJB6 LGMDD1 G/F domain mutations in the essential yeast DNAJ protein Sis1 (DNAJB6-F93L (Sis1-F106L), DNAJB6-N95L (Sis1-N108L), DNAJB6-D98Δ (Sis1-D110Δ), and DNAJB6-F100I (Sis1-F115I)). Our goal is to accelerate our understanding of mutant DNAJB6 dysfunction in LGMDD1 and thus facilitate therapeutic target identification as well as to gain insight into the relationship between protein quality control and myopathy.

The Hsp70/DNAJ machinery is vital to the protein quality control network. The Hsp70 machine works in an ATP-dependent manner to act on client proteins through a cycle of regulated binding and release^13^. Client specificity of the Hsp70 machine is modulated by DNAJ proteins (Hsp40s). By dictating client specificity, DNAJ proteins have been described as the primary facilitators of the cellular protein quality control system and play a pivotal role in determining the fate of a misfolded protein – whether is refolded or degraded^13^. A variety of disease-associated misfolded proteins have been shown to interact or colocalize with DNAJ family members^8,14^. These effects are generally, although not exclusively, dependent upon cooperation with Hsp70 ^8,13^. Strikingly, previous work from our lab suggests that the LGMDD1 mutants not only show substrate specificity, but also show conformation-specific effects ^12,15^. These results were obtained through analysis of LGMDD1 mutants in the DNAJ protein Sis1, which has well-known yeast prion protein clients. Unlike mammalian prions, yeast prions are non-toxic, but phenotypic and biochemical assays developed enable rapid detection of [*PRION+*] cells^16^. Yeast chaperones Hsp104, Hsp70 (Ssa1), and the Hsp40 (Sis1) regulate prion propagation by acting on prion protein aggregates^17^. Alterations in chaperone level or function result in a failure to promote prion propagation^18,19^.

One of the most interesting features of prions is the existence of prion strains. Prion strains are distinct self-propagating protein aggregate structures that cause changes in transmissibility and disease pathology with the same aggregating protein^20^. Yeast prion strains differ from each other based on phenotype, the ratio of soluble to aggregated protein, and their ability to propagate the prion^20^. Previously, we found that homologous LGMDD1 mutations in Sis1 appear to reduce functionality, as determined by changes in their ability to modulate the aggregated state of select yeast prion strains^12^. We then assessed the effect of these mutants in mammalian systems, including mouse models, and LGMDD1 patient fibroblasts, where we analyzed the aggregation of TDP-43, an RNA binding protein with a prion-like domain that is a marker of degenerative disease including LGMDD1. DNAJB6 mutant expression enhanced the aggregation and impaired the dissolution of nuclear stress granules containing TDP-43 following heat shock^12^. Recently, three novel pathogenic mutations associated with aberrant chaperone function that leads to LGMDD1 have been identified within the J domain of DNAJB6^21^. Interestingly, we found that homologous mutations in the Sis1 J domain differentially alter the processing of specific yeast prion strains, as well as a non-prion substrate^15^.

Here, we used the yeast prion model system, and *in vitro* chaperone activity assays to determine how LGMDD1 homologous missense mutations in the Sis1 G/F domain alter chaperone function with and without Hsp70. We found that LGMDD1 mutants inhibit the Hsp70 ATPase cycle function in a client-conformer specific manner. Moreover, both prion propagation and luciferase refolding activity were enhanced in mutant strains by either deleting the NEF (Sse1) or by using Sse1-mutants, indicating that fine-tuning of substrate processing can rescue the mutant defects. To our knowledge, this is the first mechanistic insight describing the effect of LGMDD1 mutants on Hsp70/40 ATPase cycle. Additionally, these results suggest that the development of a titrated approach using specific inhibitors of the Hsp70/DNAJ cycle is a potential therapeutic strategy for this class of myopathy-associated chaperonopathies.

## Results

### 1. LGMDD1-associated homologous G/F domain mutants in Sis1 have variable substrate processing efficiency

In previous studies, we analyzed the functionality of LGMDD1 mutants in Sis1 using two yeast prion proteins that require Sis1 for propagation: Rnq1 (which forms the [*RNQ*+] prion) and Sup35 (which forms the [*PSI*+] prion). We found that LGMDD1 mutants had conformer-specific processing defects, in that they promoted the propagation of some prion strains but not all^12^. This suggests that the mutant chaperones may recognize their normal clients in some conformations but not others. Alternatively, the mutant chaperones may recognize clients but not be able to effectively process the misfolded protein. Hsp40 client processing for refolding requires multiple steps: binding, dimerization, binding to Hsp70, and nucleotide exchange-stimulated client release. We evaluated the function of LGMDD1 mutants as compared to wild type (WT), using assays that address these steps.

One way to determine substrate processing of a chaperone that interacts with amyloidogenic proteins is to evaluate the kinetics of aggregation of client proteins *in vitro*. We assessed amyloid formation of Rnq1 in the presence of Sis1-WT and LGMDD1 mutants (G/F domain mutants Sis1-F106L, Sis1-N108L, Sis1-D110Δ and, Sis1-F115I). Rnq1 is a well-known substrate of Sis1 and has been reported to form higher-order aggregates *in vitro*^23^. Aggregation of Rnq1 was monitored by the enhanced fluorescence emission of the dye Thioflavin T (ThT), which is used as a marker for cross β sheet conformation of amyloid fibrils^24^. During the initial phase of incubation (lag phase), natively disordered protein monomer did not show any change in fluorescence intensity of ThT (Fig. 1A). This was followed by an increase in ThT fluorescence intensity, indicative of the formation of fibrils. Finally, the fluorescence intensity of the dye achieved a plateau (stationary phase). The lag phase was increased in the presence of Sis1-WT as compared to Rnq1 without chaperones (Fig. 1A), indicating that the time required for productive nucleation of unseeded Rnq1 was longer in the presence of chaperone. However, the lag phase of Rnq1 fiber formation was shorter with LGMDD1 mutants than Sis1-WT but longer than that of Rnq1 alone (Fig. 1A). The overall fibril formation rate was also higher in the presence of the mutants as compared to Sis1-WT (Fig. 1A). We also performed seeded kinetic assays with Rnq1 fibers formed at 18°C, 25°C and 37°C. We found that fiber elongation was faster (in the absence of chaperone) with seeded Rnq1 at 18°C, than at 25°C or 37°C (Fig.S1A, B, and C). Similarly, in the presence of Sis1-WT, the difference in fiber elongation using Rnq1 seeds formed at the three temperatures was negligible (Fig.S1A, B, and C). This suggests that the primary impact of Sis1 on Rnq1 fiber formation is in amyloid-competent conformer formation (nucleation) in the lag phase. Formation of aggregates was also assessed by semi-denaturing agarose gel electrophoresis (SDD-AGE). Rnq1 monomer showed a soluble band with no aggregates, while fibers of Rnq1 showed higher molecular weight aggregates (Fig. S1D).

**Figure 1.**
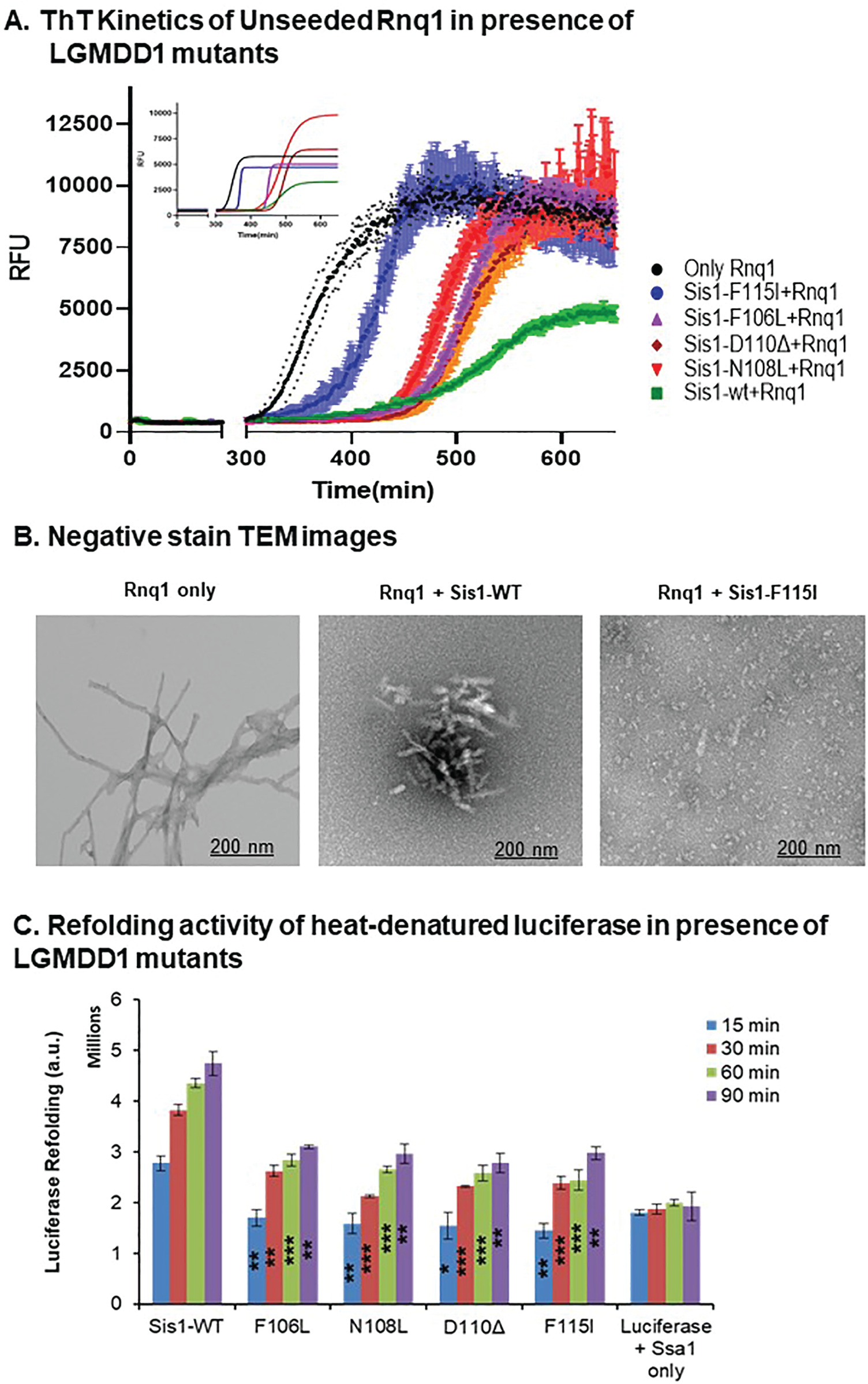
LGMDD1 G/F domain mutants show variability in substrate processing. (A) Kinetics of Rnq1 fibrillation in the presence of unseeded Rnq1 only **(black)**, Sis1-WT **(green)**, Sis1-F106L (**purple**), Sis1-N108L (**red**), Sis1-D110Δ (**brown**), or Sis1-F115I (**blue**) measured by ThT fluorescence assay. *Inset:-* Fitted graph using the aggregation kinetics equation *y*=y*i* +mxi +(yf+mxf)/1+(ex ^χ0/τ^) where (yi + mxi) is the initial line, (yf + mxf) is the final line, and x0 is the midpoint of the maximum signal. (B) The morphology of amyloid fibers formed from Rnq1 at 18°C in vitro in the presence and absence of Sis1-WT and/or LGMDD1 mutant were imaged by TEM. The scale bar represents 200 nm. (C) Refolding activity of heat-denatured luciferase in the presence of LGMDD1 mutants. Luciferase along with Ssa1-WT and Sis1-WT/mutants was incubated at 42°C for 10 min to heat denature luciferase. At various time points, activity was measured by a luminometer after adding substrate. Each LGMDD1 mutant was compared with Sis1-WT across all time-points. ***p<0.001, **p<0.01, and *p<0.05.

We then hypothesized that, if the primary chaperone effect is to alter the nucleation of the amyloidogenic protein, the fibers formed with and without chaperone could be structurally distinct. As such, we assessed the morphology of Rnq1 fibrils formed at 18°C with and without chaperone by Transmission Electron Microscopy (TEM). TEM images show that Rnq1 forms long, elongated, branched fibrils in the absence of chaperone (Fig. 1B). By contrast, Rnq1 fibers formed in the presence of Sis1-WT were short and appeared immature or perhaps bound to chaperone (Fig. 1B). This suggests that Sis1-WT might only delay fibril formation rather than preventing it. In the presence of LGMDD1 mutant (Sis1-F115I), very few fibers were visible (Fig. 1B) and they were mostly distorted and appeared to be small oligomeric species. These data suggest that the interaction of LGMDD1 mutants with client change the conformation of client in a manner that is distinct from that of wildtype chaperone.

While the interaction with prion substrate showed that LGMDD1 mutants alter amyloid formation, DNAJ proteins typically act in conjunction with Hsp70 to process substrates. We then asked whether the LGMDD1 mutants could promote substrate refolding in the presence of co-chaperones. Previously Aron et. al. showed that Sis1 lacking the domain that harbors these LGMDD1 mutants (Sis1ΔG/F) is defective in chaperone activity and partially inhibits the ability of Sis1-WT to facilitate folding of denatured luciferase protein^25^. Therefore, to test the ability of LGMDD1 mutants to function in substrate refolding, we heat-denatured firefly luciferase and then monitored refolding in the presence or absence of chaperones. We found that the LGMDD1 mutants were all compromised in their ability to refold luciferase (Fig. 1C). Taken together, these results indicate that the LGMDD1 mutants are defective in substrate processing.

### 2. LGMDD1-associated G/F domain mutants change the formation of *in vitro* formed [*RNQ*+] prion strains

Given the striking differences in fibril formation kinetics and TEM images, we wanted to determine whether LGMDD1 mutants alter the formation of Rnq1 amyloid structures using a more sensitive and quantitative method that is uniquely available in the yeast prion system. This entails infecting yeast with amyloid generated in vitro and assessing the resulting [*RNQ+*] prion strains. Previously, we found that amyloid fibers of Rnq1PFD (prion-forming domain) formed at different temperatures resulted in the generation of different prion strains^26^.

We hypothesized that Rnq1 amyloid structure, and resulting [*RNQ+*] strain distribution, would differ in the presence of Sis1 and LGMDD1 mutants. We formed Rnq1 fibers at 18°C in the presence and absence of Sis1-WT and two LGMDD1 mutants. We transformed fibers formed in the presence or absence of chaperone into cells expressing [*RNQ*+] reporter protein (RRP) that did not have the prion ([*rnq*-]) (Fig. S2A). We developed and utilized this reporter strain as a phenotypic indicator of [*RNQ*+] prion strain propagation^26^. By assessing colony color (white, light pink, or dark pink) as well as growth on selective medium (SD-Ade), we scored [*RNQ+*] strains (Fig. 2A and B) that resulted after fiber infection. Lighter colony color and more growth on SD-Ade medium was scored as a stronger [*RNQ*+] strain, whereas darker colony color and less growth on selective medium was scored as a weaker [*RNQ*+] strain (see Methods for more information). [*RNQ+*] strains were also verified for the prion-specific trait of curability on medium containing guanidine hydrochloride (Fig. 2A and B). To quantify the difference in [*RNQ+*] strains formed in the presence of LGMDD1 mutants we counted the phenotype of infected colonies from five different transformation sets for each sample (Fig. 2C). With Rnq1 alone, ~58% of cells showed very weak [*RNQ+*] phenotypes. This fraction of very weak [*RNQ+*] was reduced to less than 20% in the presence of Sis1-WT and was further reduced in the presence of the LGMDD1 mutants. There was a significant increase in the medium [*RNQ+*] strain phenotype (44% +/− 4.5%) in the presence of Sis1-WT as compared to Rnq1 alone (14% +/− 1.4%). When we compared [*RNQ+*] strain distribution between Sis1-WT and LGMDD1 mutants, we found that the proportion of weak Rnq1 strains was significantly increased with both F106L and F115I (~20% in Sis1-WT vs. 50% in F106L and 65% in F115I). Similarly, the population of medium [*RNQ+*] strain was significantly decreased in F115I as compared to Sis1-WT (~20% in F115I vs. 45% in Sis1-WT). The distribution of other [*RNQ+*] strains, strong and very weak, in LGMDD1 mutants were similar to Sis1-WT. Taken together, the difference in distribution of [*RNQ+*] strains in the presence of LGMDD1 mutants establish both the conformer-specific effects and suggest that the LGMDD1 mutants are not simply loss-of-function mutants as the [*RNQ*+] strains generated are markedly different than those arising in the absence of chaperone.

**Figure 2.**
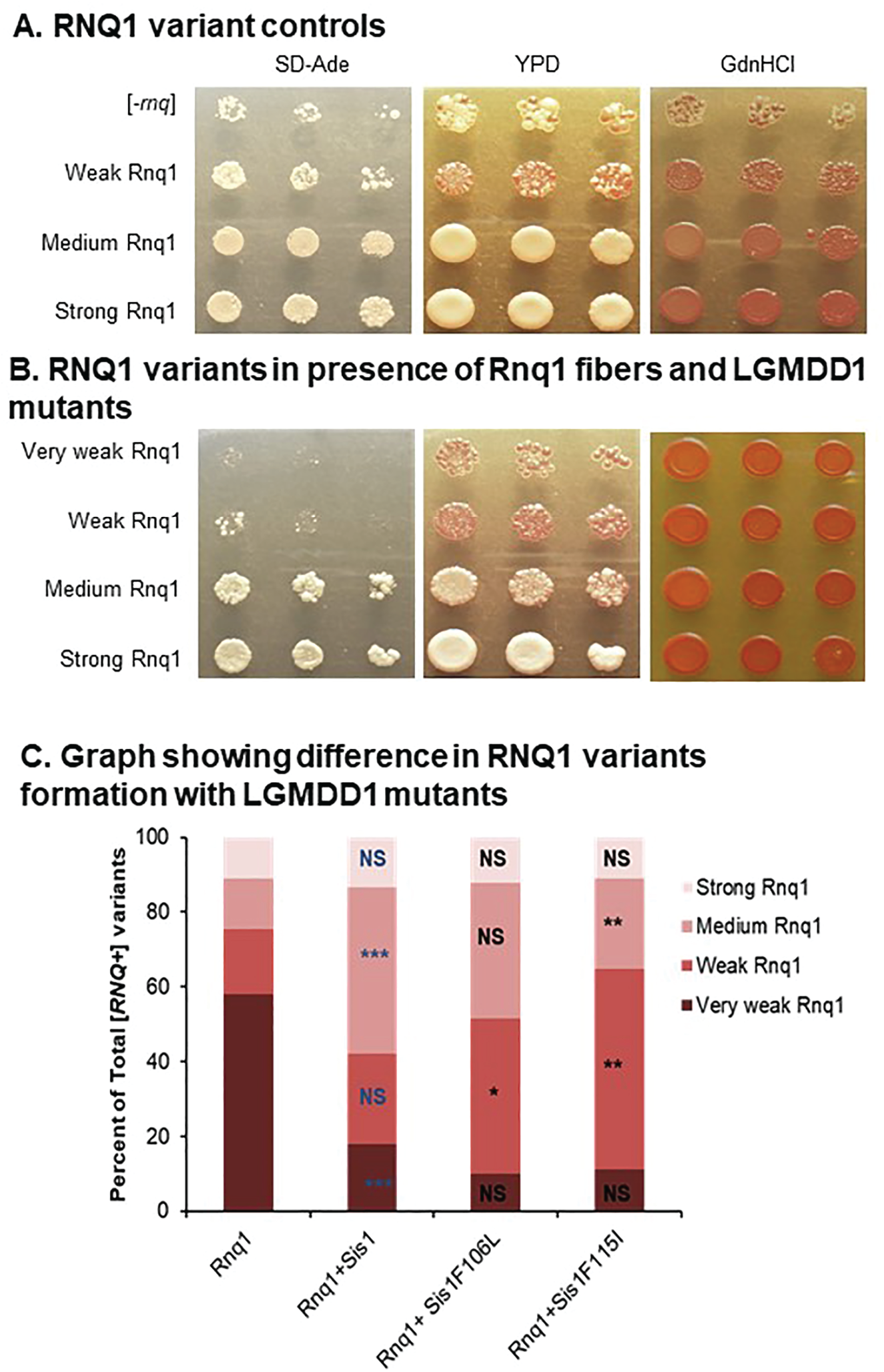
Phenotypic distribution of [*RNQ+*] strain alters with LGMDD1 G/F domain mutants. Transformation of Rnq1 amyloid fibers formed at 18°C into [*rnq-*] 74D-694 yeast cells induces strains of the [*RNQ+*] prion (A). RRP was used to assess [*RNQ+*] phenotype. [*RNQ+*] prion strains co-aggregate with RRP and cause different phenotypes; colony color on YPD and growth on SD-ade medium are indicative of different levels of suppression of the *ade1-14* premature stop codon to produce Ade1 and represent different [*RNQ+*] strains. Curability by growth and color on medium containing GdnHCl was used to determine prion-dependence of phenotypes. The transformants from five separate experiments for each sample set were picked and >200 colonies for each set were scored as very weak, weak, medium, or strong [*RNQ+*] (B) and graphed for statistical analysis (C). In panel (C), the blue color indicates the comparison between Rnq1 (alone) and Rnq1 with Sis1-WT, black color indicates the comparison between Sis1-WT and LGMDD1 mutants (F106L and F115I). ***p<0.001, **p<0.01, *p<0.05, and NS is non-significant.

### 3. LGMDD1-associated homologous mutants in Sis1 (Hsp40) alters its ability to function with Ssa1 (Hsp70) efficiently

Hsp40s target substrates to their Hsp70 partners and regulate the ATPase activity and substrate binding of the Hsp70^13^. The recognition of substrates depends on their conformation, and it has been suggested that much of the Hsp40 conformation-dependent recognition is dependent on the G/F domain ^22,27–29^. Thus, we evaluated the ability of LGMDD1 mutants to function in conjunction with Ssa1. We measured the ability of Sis1-WT and LGMDD1 mutants to stimulate the ATPase activity of Ssa1 in the presence and absence of different Rnq1 protein conformers. We standardized the Rnq1 monomer concentration and performed a phosphate standard curve with each assay (Fig. S3A). Notably, there was no difference in ATP hydrolysis rate between Sis1-WT and LGMDD1 mutants in the absence of any client protein (Fig. S3B). However, the ATP hydrolysis rate was significantly reduced with the mutants in the presence of Rnq1 seeds formed at 18°C and 25°C (Fig. 3A and B). Interestingly, the ATP hydrolysis rate of all mutants was marginally but not significantly reduced as compared to Sis1-WT in the presence of Rnq1 seeds formed at 37°C (Fig. 3C). We found no difference in ATP hydrolysis rate between Sis1-WT and LGMDD1 mutants in the presence of Rnq1 monomer (denatured) as client (Fig. 3D). This may be due to the fact that “denatured monomer” presents a variety of epitopes that can be recognized by chaperone whereas the fibers likely have fewer sites for recognition. These results indicate that the LGMDD1 mutants’ ability to stimulate the ATPase activity of Ssa1 is conformer-dependent.

**Figure 3.**
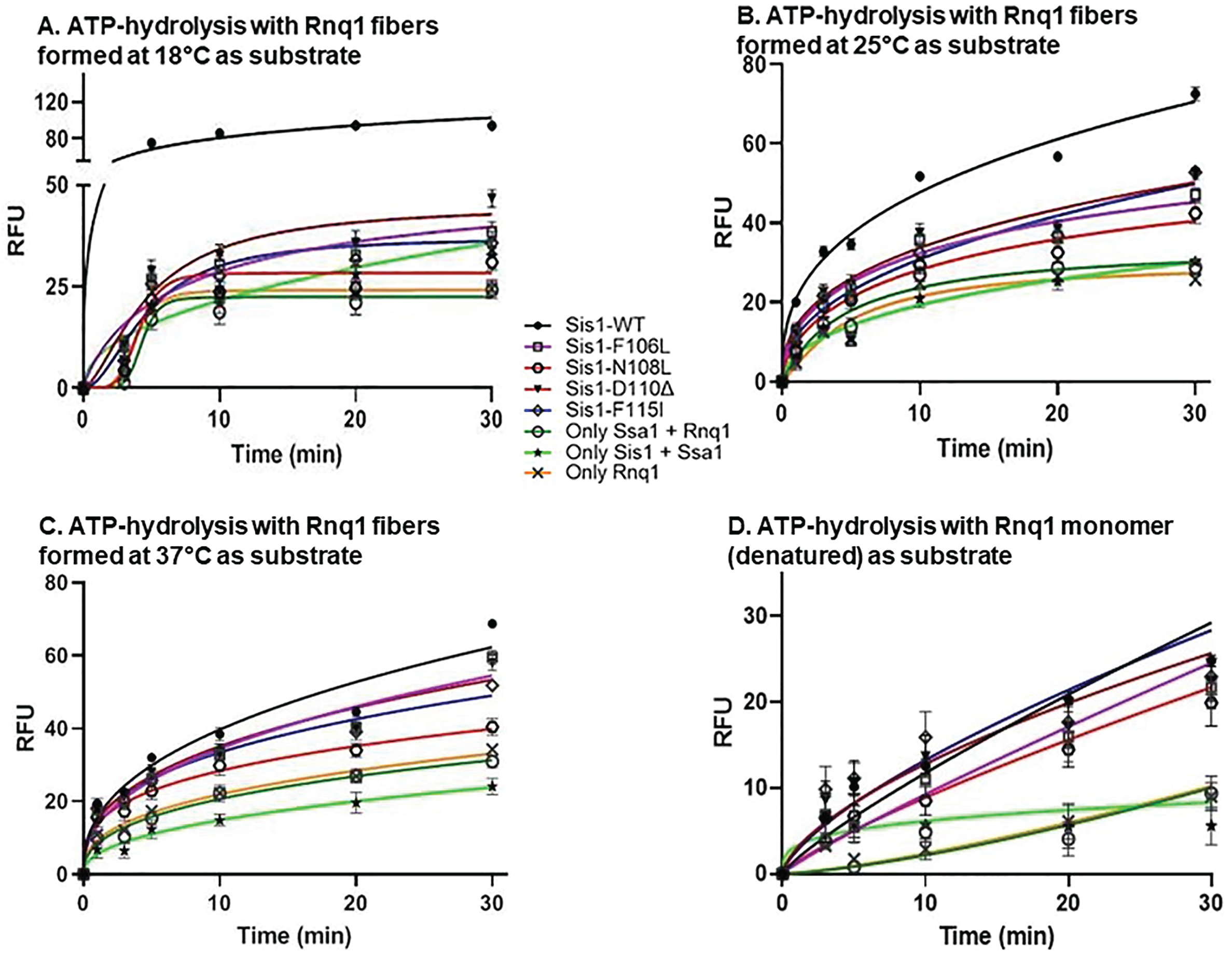
Stimulation of ATPase activity of Ssa1 by LGMDD1 G/F domain mutants is client-conformation specific. Stimulation of Ssa1 ATPase activity in the presence of Rnq1 seeds formed at (A) 18°C, (B) 25°C, and (C) 37°C, and (D) Rnq1 monomer. Ssa1 (1 μM) in complex with ATP (1 mM) in the presence of Sis1-WT (**black**) or Sis1-mutants (Sis1-F106L (**purple**), Sis1-N108L (**red**), Sis1-D110Δ (**brown**), or Sis1-F115I (**blue**)) (0.05 μM). The fraction of ATP converted to ADP was determined at indicated times. For (A), (B), and (C) a total of 10% seeds were used in a reaction. For (D) Rnq1 monomer used was 25 μM. In all cases, only Rnq1 (**orange**), Sis1 with Ssa1 (**light green**) and Ssa1 with Rnq1 (**dark green**) were used as controls. For A-D, Sis1-WT was compared with LGMDD1-mutants at all time points. Here we show the significant difference at one time-point (20 min). For (A) and (B) values were ***p<0.001, for (C) values were **p<0.01, for (D) values were non-significant (NS).

### 4. LGMDD1 G/F domain mutants alters both substrate and Hsp70 binding

There are multiple facets that determine the efficiency of Hsp70-40 ATPase cycle. One of the key initial steps is the ability of Hsp40 to bind to client. Thus, we wanted to test the physical and functional interactions of LGMDD1 mutants with our Rnq1 and luciferase substrates. We performed a binding assay utilizing a method previously used to study interactions between the *E. coli* DnaJ and substrates^30^. We found that LGMDD1 mutants show significantly reduced binding to denatured Rnq1 and luciferase substrates as compared to Sis1-WT (Fig. 4A and B) at low concentrations of substrate. Once the mutants bind substrate, however, they are able to stimulate Hsp70 activity (Fig. 3D).

**Figure 4.**
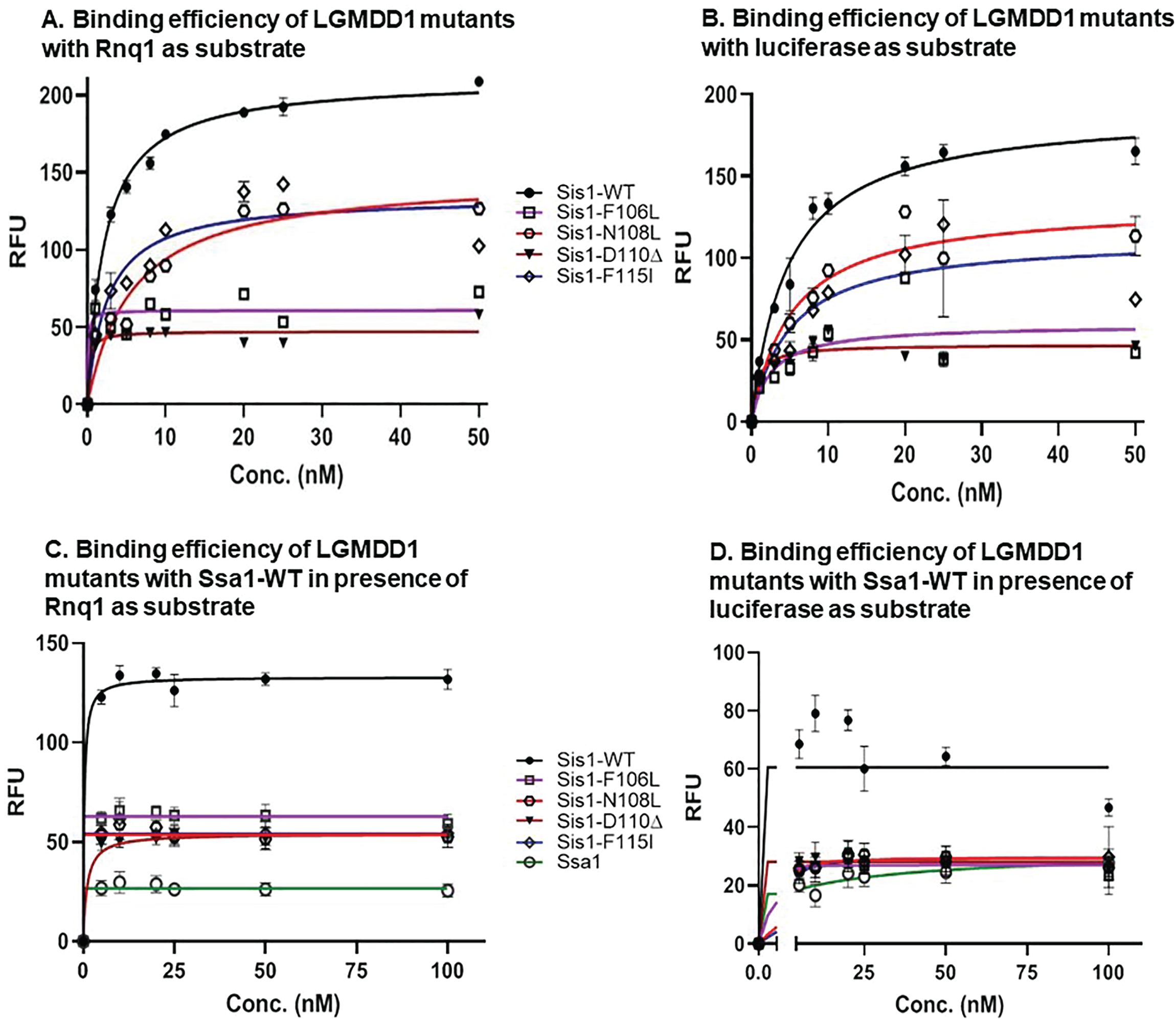
Sis1 binding to both substrate and Hsp70 is compromised in the presence of LGMDD1 G/F domain mutants. Binding of purified Sis1-WT (**black**), Sis1-F106L (**purple**), Sis1-N108L (**red**), Sis1-D110Δ (**brown**), or Sis1-F115I (**blue**) to denatured Rnq1 (A) and luciferase (B) Rnq1 (400 ng) and luciferase (100 ng) immobilized in microtiter plate wells and dilutions of purified Sis1-WT and Sis1-mutants (0, 1, 3, 5, 8, 10, 20, 25, 50 nM) were incubated with each substrate. The amount of Sis1 retained in the wells after extensive washings was detected using a Sis1 specific antibody. Denatured Rnq1 (C) and denatured luciferase (D) were premixed with Sis1-WT/mutants and immobilized in microtiter plate wells and dilutions of Ssa1-WT (0-100 nM) were incubated with it. Bound Ssa1-WT was detected using an αSsa1 antibody. For A-D, Sis1-WT was compared to LGMDD1 mutants. For (A-D), values shown are ***p<0.001.

A key function for Hsp40s is stimulating Hsp70s. Based on our previous work, we hypothesized that the deleterious effect of the DNAJB6 G/F domain mutants is HSP70-dependent^31^. Thus, the LGMDD1 mutants might be altered in their productive association with Hsp70. These mutants may affect the cycle by either reducing Hsp70 binding, sequestering Hsp70, or they may be hyperactive and alter the refolding process. We found that reducing the interaction of LGMDD1 mutants with Hsp70 using pharmacologic compounds led to improvement in muscle strength and myopathology in mouse models^31^. Therefore, we performed binding assays to determine whether there was a productive association between the LGMDD1 mutants and Ssa1. We found that the interaction between LGMDD1 mutants and Ssa1 was significantly reduced both in the absence (Fig. S4) and presence of client proteins Rnq1 (Fig. 4C) and luciferase (Fig. 4D). This indicates that the reduced interaction between the mutants and Ssa1 is client-independent.

### 5. LGMDD1 G/F domain mutants show reduced dimerization efficiency

The efficient function of Sis1 requires the ability of the protein to form dimers. In fact, Sis1 (1– 337aa), which lacks the dimerization motif, exhibited severe defects in chaperone activity, but could regulate Hsp70 ATPase activity^32^. Sha et. al. proposed that the Sis1 cleft formed in dimers functions as a docking site for the Hsp70 peptide-binding domain and that this interaction facilitates the transfer of peptides from Sis1 to Hsp70^33^. Thus, in order to further evaluate the chaperone activity of the LGMDD1 mutants, we set out to measure their dimerization efficiency. We performed binding assays to determine the relative competence of both the self-association of the mutants as well as their association with Sis1-WT. We found that there was a significant decrease in self-association of all the mutants as compared to Sis1-WT (Fig. 5A). We also found that their ability to bind to Sis1-WT was significantly reduced (Fig. 5B), indicating a possible explanation for the reduction in chaperone activity observed with these mutants.

**Figure 5.**
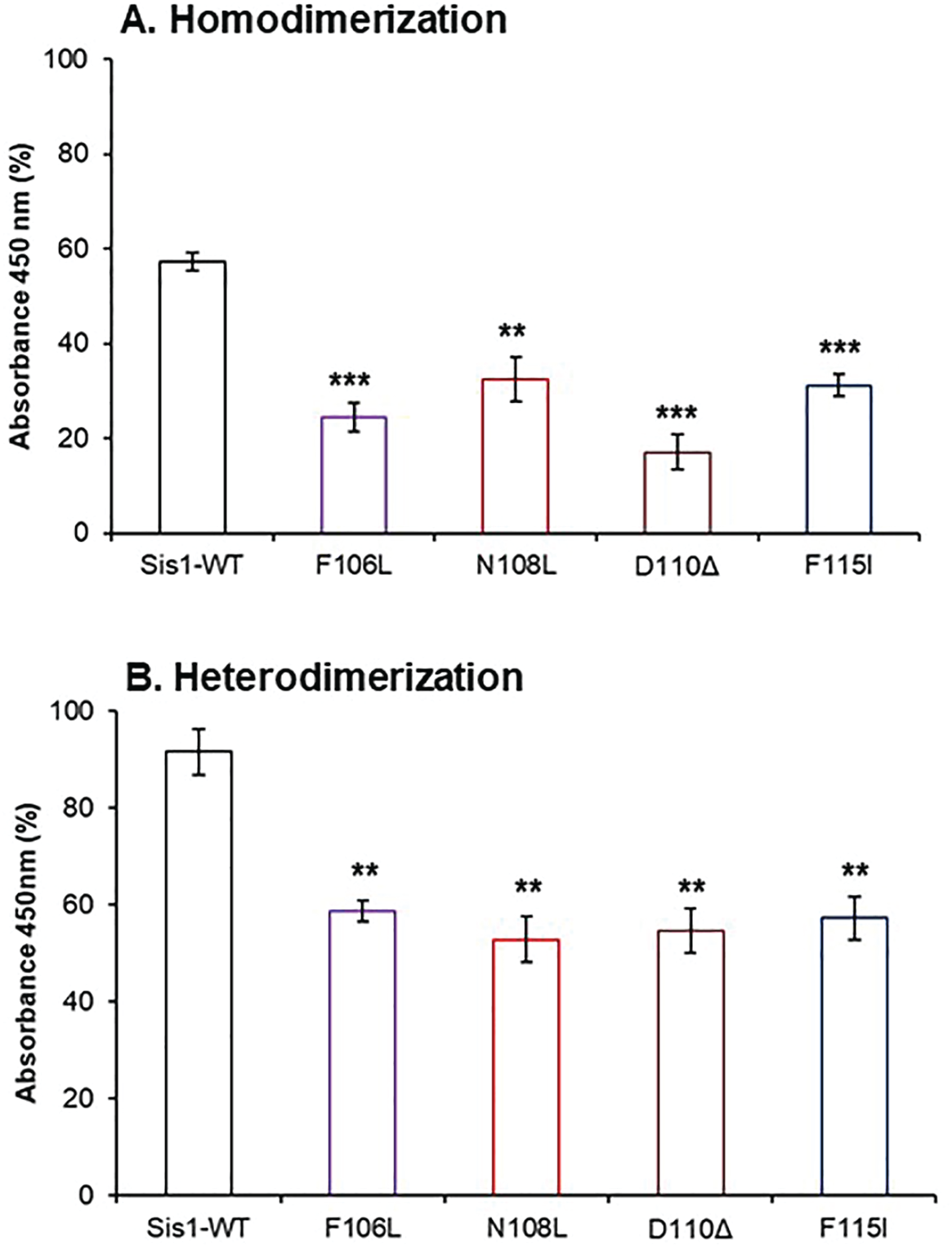
LGMDD1 G/F domain mutants were defective in dimerizing. (A) For homodimerization, uncleaved His-tagged Sis1-WT and mutants (20 nM) were added into non-His-tagged (cleaved) Sis1-WT and mutants (200 nM) and adsorbed in microtiter plate wells. (B) For heterodimerization, uncleaved His-tagged Sis1-WT (20 nM) was added to cleaved mutants (200 nM) and adsorbed in microtiter plate wells. In both (A) and (B), following adsorption, ELISA was performed using an anti-His antibody. All LGMDD1 mutants were compared with Sis1-WT.

### 6. LGMDD1 G/F domain mutants inhibit Sis1-WT induced Ssa1 ATPase activity by reducing its dimerization and substrate binding efficiency

Due to the dominant nature of these mutants, we set out to test the effect of LGMDD1 mutants on the Sis1-WT induced ATPase activity of Ssa1 in the presence of different client-conformers. For this, we performed an ATPase assay with mixtures of Sis1-WT and LGMDD1 mutants; F106L and F115I in the presence of Rnq1 fibers formed at two different temperatures, 18°C and 25°C. We found that there was a gradual decrease in the rate of ATP hydrolysis that correlated with titrating Sis1-WT with F106L (Fig. 6A and S5A) and F115I (Fig. 6B and S5B).

**Figure 6.**
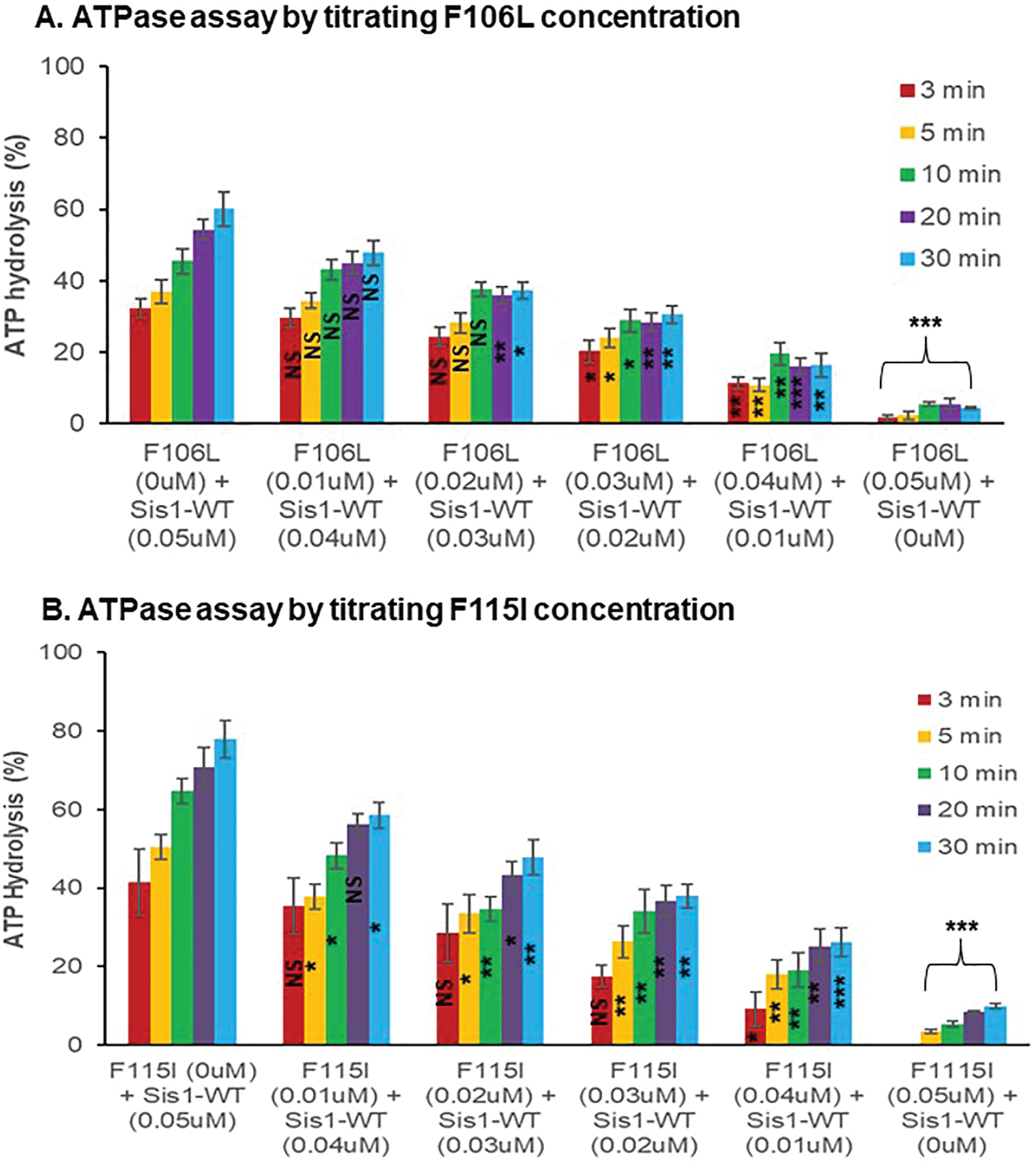
LGMDD1 G/F domain mutants inhibit Sis1-WT induced ATPase activity of Ssa1. Stimulation of Ssa1 ATPase activity in the presence of Rnq1 seeds formed at 18°C. Sis1-mutants (Sis1-F106L (**A)**, or Sis1-F115I (**B**)) (0-0.05 μM) were titrated with Sis1-WT (0-0.05 μM) in the presence of Ssa1 (1 μM) and ATP (1 mM). The fraction of ATP converted to ADP was determined at various time-points from 3 to 30 minutes. Values of the increasing concentration of LGMDD1 mutants (0.01-0.05 μM) were compared with Sis1-WT (0.05 μM) alone.

Efficient ATP hydrolysis is a culmination of many small but significant events related to the individual function of Sis1 (Hsp40), Ssa1 (Hsp70) and other players in the ATP-hydrolysis cycle. Two critical aspects of Sis1-WT activity are its ability to dimerize and to bind to substrate efficiently. LGMDD1 mutants show decreased dimerization as compared to wild-type protein (Fig 5A and B). Thus, we decided to determine whether the mutants also inhibit the ability of wild-type protein to dimerize. We found that there was a concomitant decrease in dimerization with the titration of Sis1-WT with F106L (Fig. S6A) and F115I (Fig. S6B). We asked whether the mutants also inhibit the ability of wild-type protein to bind substrates efficiently. The binding efficiency to both substrates, Rnq1 (Fig. S6C) and luciferase (Fig. S6D), were slightly but significantly reduced when Sis1-WT was used in equal proportion (1:1) with each of the LGMDD1 mutants F106L and F115I. These data provide mechanistic insight into the inhibition of Sis1-WT-induced Ssa1 ATPase activity in the presence of LGMDD1 mutants (Fig. 6).

### 7. Modulating Hsp40-Hsp70 cycle by either deleting or inhibiting nucleotide exchange factors (NEFs) can be beneficial with respect to LGMDD1 G/F mutant effect *in vivo*

The lifetime of the Hsp70/40:substrate complex is dependent upon nucleotide exchange. A key player in this is nucleotide exchange factors (NEFs) that stimulate ADP release. In yeast, cytosolic Hsp70 interacts with three NEFs homologous to human counterparts: Sse1/Sse2 (Hsp110), Fes1 (HspBP1), and Snl1 (Bag-1)^34,35^. Previously, we found that LGMDD1 mutants impair viability and prion propagation in yeast and these effects were rescued by reducing the association with Hsp70^31^. Thus, we decided to investigate whether deleting Hsp110 (Sse1) would have a similar rescuing effect. A significant effect would further support the hypothesis that Hsp70 activity inhibition provides a potential mechanism for therapeutic intervention for LGMDD1-associated myopathy. To assess the impact of NEFs, we asked whether their deletion would enhance the ability of LGMDD1 mutants to propagate [*RNQ*+] prion strains. We used two established biochemical yeast prion assays that differentiate soluble and aggregated protein: well-trap and boiled gel assays. We found that the deletion of Sse1 partially rescues [*RNQ*+] prion propagation (Fig. 7A, S7A). However, it was specific to Sse1, as the alteration of the other NEFs did not demonstrate such rescuing (Fig. S7B). This was not surprising, as Sse1 is the principal NEF in yeast and performs 90% of the NEF activity in the cell^36^.

**Figure 7.**
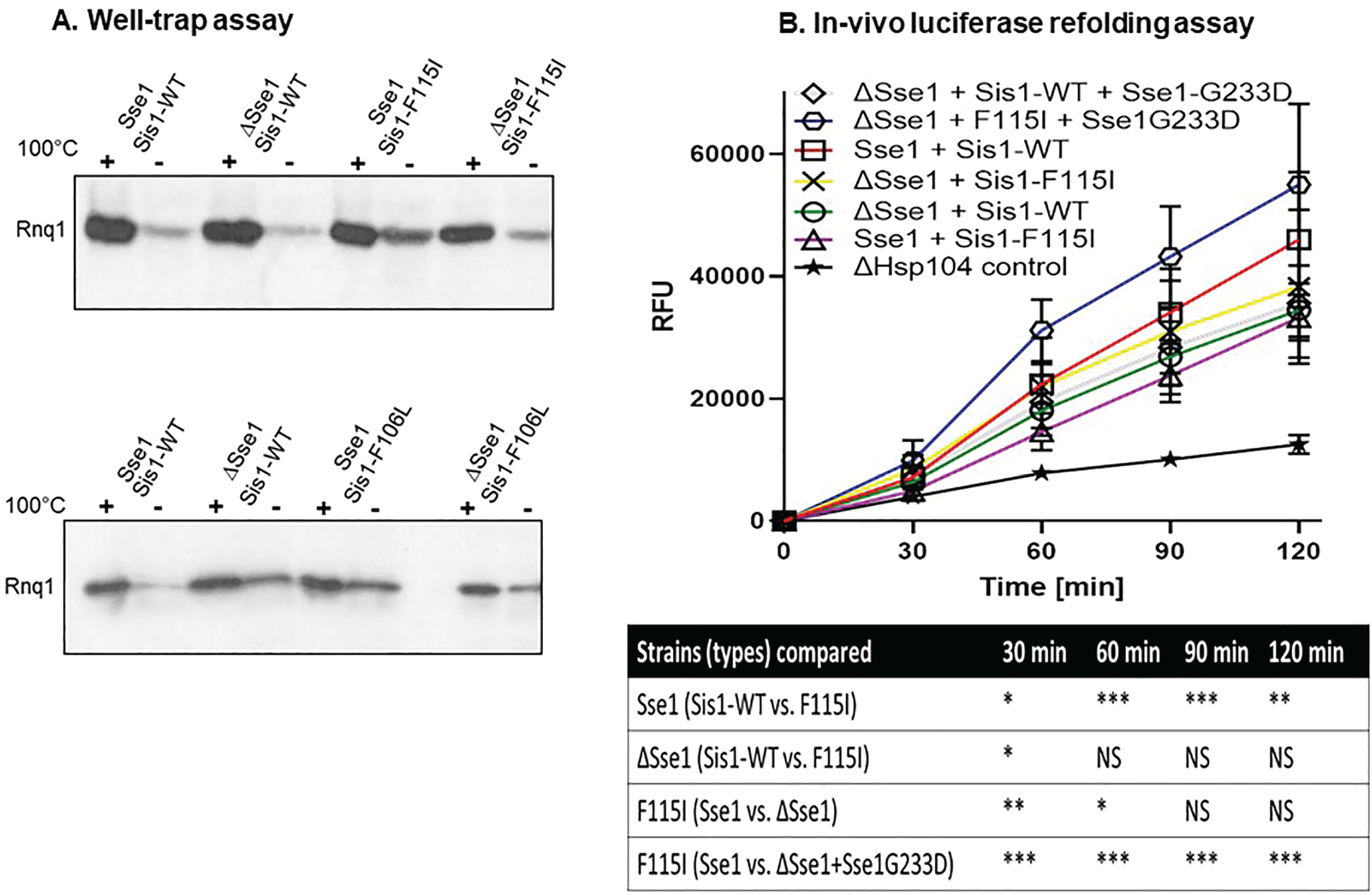
Deletion or inhibition of Sse1 function rescues prion propagation of [*RNQ+*] and restored impaired FFL refolding. (A) Deletion of Sse1 partially suppresses prion loss caused by LGMDD1 mutants. Analysis of prion propagation in Sse1 and *Δsse1* strains harboring s. d. medium [*RNQ+*], expressing either Sis1-WT or Sis1-F115I or Sis1-F106L by well-trap assay. Cell lysates were incubated at room temperature (-) or 100°C and subjected to SDS-PAGE and western blot using an αRnq1 antibody. Rnq1 that is not sequestered in aggregates will enter the gel in the unboiled sample, which indicates destabilization of the [*RNQ+*] prion. (B) The refolding of firefly luciferase (FFL) was measured in the Sse1 and *Δsse1* yeast cells carrying Sis1-WT, LGMDD1 mutant (Sis1-F115I), and Sse1 mutant (Sse1-G233D) along with a plasmid expressing FFL. Yeast were normalized, treated with cycloheximide, and subjected to heat shock at 42°C for 22 minutes, followed by recovery at 30°C. Luminescence was measured at the indicated time points during recovery. Values shown are ± SEM of at least three independent experiments.

Based on these data, we hypothesized that fine-tuning the inhibition of Hsp70 activity could have a positive phenotypic effect with respect to LGMDD1 mutants. Thus, we assessed the effect of characterized Sse1 mutants that inhibit Hsp70 activity either by delaying binding to the Hsp70-client-ADP complex (Sse1-K69Q) or by delaying the release of ADP from the Hsp70-client complex (Sse1-G233D)^37^. We performed boiled gel assays to examine the relative levels of soluble Rnq1 prion protein in [RNQ+] cells. Indeed, we found a decrease in soluble Rnq1 protein (like in Sse1-WT cells expressing Sis1-WT) in LGMDD1 cells expressing either Sse1-G233D (Fig. S7C) or Sse1-K69Q (Fig. S7D). This indicates restoration of [*RNQ+*] prion propagation in LGMDD1 mutants with NEF modulation. This further supports our hypothesis that fine-tuning Hsp70 activity could provide a therapeutic avenue for LGMDD1.

To further understand the impact of Sse1 on the LGMDD1 disease-associated mutants, we assessed the refolding of a non-prion substrate (firefly luciferase; FFL). We utilized an *in vivo* refolding assay in which FFL is denatured in cells by heat shock and its subsequent refolding, which requires the Hsp40/Hsp70/Hsp104 chaperone machinery, is measured by activity^38,39^. We transformed the Sse1 and *Δsse1* yeast cells carrying Sis1-WT, LGMDD1 mutants, and Sse1 mutants with GPD-lux vector for the expression of FFL. Since Hsp104 is required for efficient refolding of FFL, we used a *Δhsp104* strain expressing FFL as a negative control. We found that the FFL refolding activity was significantly altered in Sse1 cells expressing F115I as compared to cells expressing Sis1-WT (Fig. 7B). This difference in FFL refolding activity between LGMDD1 mutants (F115I) and Sis1-WT was non-existent in *Δsse1* yeast cells (Fig. 7B). However, we observed that the FFL refolding activity was marginally higher in *Δsse1* yeast cells expressing F115I as compared to Sse1-WT cells expressing the same mutant protein (Fig. 7B). Interestingly, *Δsse1* yeast cells co-expressing F115I and Sse1 mutant (Sse1-G233D) showed significant improvement in FFL refolding activity as compared to Sse1-WT yeast cells (Fig. 7B). This improvement in FFL refolding activity with Sse1-mutants correlated with the improvement of Rnq1 prion propagation observed in *Δsse1* yeast cells (Fig. S7C and S7D). As cellular homeostasis depends upon the efficient functioning of the entire chaperone machinery, our results indicate that LGMDD1 mutants delay Hsp70 ATPase activity, possibly resulting in the increased load of aggregation-prone muscular proteins observed in LGMDD1 patients.

## Discussion

Proper functioning of protein chaperones, including that of Hsp70/DNAJ, is important for the maintenance of muscle function^40^ (Fig. 8A). Previously, we have shown that the LGMDD1 mutations in the G/F domain of DNAJB6 disrupt client processing in both a substrate- and conformation-specific manner^12^. Here, we delved into the mechanism underlying the defect in client processing shown by LGMDD1 G/F domain mutants. Our data show that the LGMDD1 mutants inhibit the Hsp70 ATPase activity (Fig. 8B), which may result in general impairment of protein quality control and accumulation of protein inclusions in muscle.

**Figure 8.**
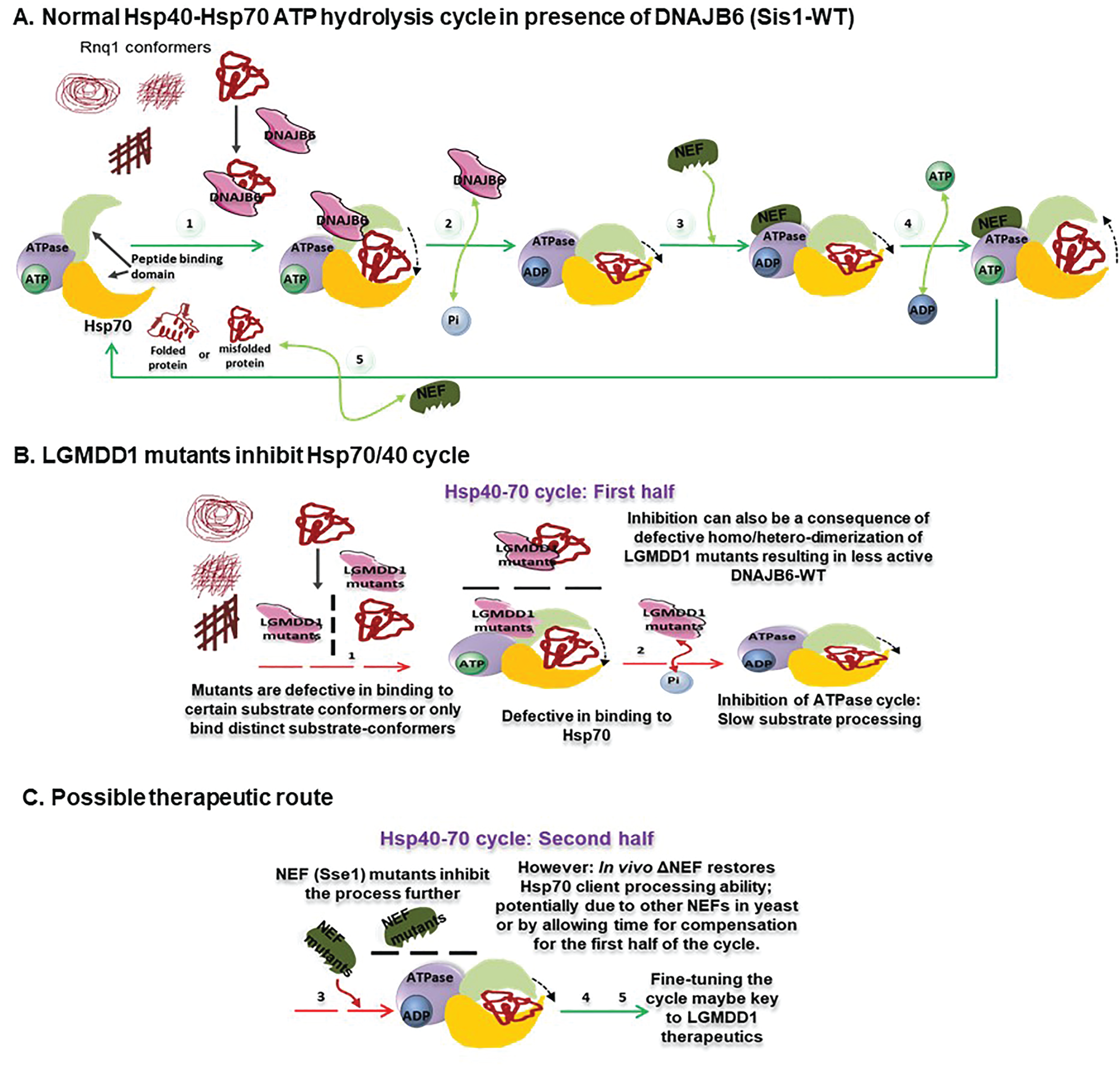
Schematic diagram depicting possible mechanism of LGMDD1 mutants and their effects on the Hsp70 ATPase cycle, as well as proposed therapeutic intervention. (A) Cartoon depicting the normal processing of one of the conformers (substrate) of an aggregated protein, mediated by DNAJB6 (Hsp40) through the Hsp70 ATPase cycle. The **green** arrows throughout the cycle indicate normal functioning. (B) Cartoon depicting possible mechanistic insights as to how LGMDD1 G/F domain mutants delay or inhibit the Hsp70 ATPase cycle. The dashed **red** arrows indicate inhibition. Mutants were inefficient in terms of both homo and hetero-dimerization and that may be related to their reduced ability to assist Hsp70 in protein folding. In the first half of the Hsp70 ATPase cycle, LGMDD1 mutants were incompetent in their ability to bind specific substrate conformers and Hsp70, which delays the downstream processing of the substrate through Hsp70-ATPase cycle. This inhibition could negatively impact Hsp70-mediated ATP-hydrolysis. (C) Possible therapeutic route. In the second half of the cycle, NEF mutants inhibit the binding and exchange of nucleotide, which delay the downstream process of substrate release in the cycle. However, this delay led to positive consequences in terms of yeast prion propagation. Overall, altering the balance between the two halves of the Hsp70-ATPase cycle may provide a key step to consider for therapeutic intervention for these types of diseases.

Indeed, we demonstrated using the yeast prion model system which affords a unique conformer:phenotype read-out. Prion proteins can form several unique prion variants (or strains) that have slight differences in their β-sheet structure that constitute distinct amyloid conformations^41^. Such different structures are presumably the underlying cause of the diverse phenotypic variation seen in both yeast and prion diseases^20^. Several mammalian pathological proteins have also been shown to adopt distinct self-propagating aggregates or “strains” with different structures, which are presumably linked to the phenotype diversities of degenerative diseases^16,42–44^. One such example is Tau protein, deposition of whose pathological forms results into Tauopathies, which includes Alzheimer’s disease, Fronto-Temporal Dementia (FTD). Several studies support the “tau strain hypothesis”, which proposes that different aggregated tau conformers (distinct strains) have distinct pathology-initiating capacities because they interact with endogenous tau differently^45,46^. In yeast, interaction between the prion protein Rnq1 and the Hsp40 Sis1 is required for [*RNQ+*] propagation^29^. Therefore, we utilized this system to ask whether the LGMDD1 mutants impacted the [*RNQ+*] prion strains that form *de novo.* Strikingly, we found a change in [*RNQ+*] prion strain formation when Rnq1 protein had formed fibers in the presence of the LGMDD1 mutants as compared to wild type Sis1. These changes were a direct consequence of Hsp40 interaction alone and may be a consequence of DNAJ proteins ability to act as *“holdases”*^47,48^. *“Holdases”* are chaperones that do not use ATP and simply protect their client protein from aggregation^48^. Unlike “*holdases*”, *“foldases”* (like Hsp70) accelerate the transition of non-native conformations towards native states in an ATP-dependent manner. Interestingly, the proteostasis network relies on a constant interplay between these two kinds of chaperones. Our results suggest that the *“holdase”* activity associated with LGMDD1 mutants is compromised and this alters substrate folding.

As a major role for Hsp40s is stimulating Hsp70s, another important aspect of the LGMDD1 mutants might be a change in the productive interaction with Hsp70^25^. Indeed, we found that these LGMDD1 mutants were defective in binding to Ssa1 (Hsp70) as well as to substrates (Rnq1 and luciferase). It had been shown previously that there was no difference in the ATPase activity between Sis1-WT and the Sis1-G/F domain knockout in absence of substrate^25^. However in our case, the introduction of the LGMDD1 mutants led to reduction in the ATPase activity of Hsp70. Moreover, in the presence of substrate, the change in ATPase activity was client-conformer specific, as assays with Rnq1 fibers formed at 18°C and 25°C showed reduced activity, while Rnq1 monomer did not. Additionally, LGMDD1 mutants were defective in refolding heat-denatured luciferase (similar to previous findings with Sis1ΔG/F), indicating a more global defect in substrate remodeling. These findings correlate well with recent data that implicate the G/F-rich region of DNAJB1 in an autoinhibitory mechanism that regulates the major class B J-domain proteins (JDPs)^49^. Thus, although in all JDPs the interaction of the J-domain is responsible for the activation of Hsp70, in DNAJB1, due to this autoinhibitive interaction with the G/F domain, the activation of Hsp70 is inhibited. This inhibition can be released with second site mutations (E50A, or F102A, and or ΔH5) on DNAJB1^49^. As DNAJB6 (Sis1) belongs to the same class B JDPs, a similar mechanism of interaction is likely, and mutations in the G/F domain could disrupt the autoinhibitory mechanism. Additionally, this is consistent with recent observation, where we have found that aggregation-independent toxicity induced by the overexpression of the Sis1 LGMDD1 F106L and F115I mutants in yeast can be rescued by reducing Hsp70 binding^31^. These results confirm that the effect of the LGMDD1 mutants is not independent of Hsp70 (at least for these substrates).

The function and interaction of the various Hsp40 domains have been studied extensively^13,50^. The efficiency of Hsp70 ATPase activity is heavily dependent on the proper functioning of the J and G/F domains of Hsp40^50,51^. Moreover, Hsp40 (Sis1) functions as a dimeric protein^32^. Interestingly, we found that LGMDD1 mutants show a diminished ability to form homodimers and heterodimers (with Sis1-WT). Thus, the LGMDD1 mutants show many defects that may contribute to perturbations in substrate processing.

Previously, it was suggested that the LGMDD1-causing mutations exert a deleterious dominant negative effect on the wild-type protein^3^. An excess of mutant (DNAJB6-F93L) to wild-type mRNA induced lethality in embryos, while an excess of wild-type to mutant mRNA gave rise to progressively increased rescue^3^. Consistent with this, we found that titrating Sis1-WT with an excess of LGMDD1 mutants (F106L and F115I) decreased ATPase activity.

Interaction of the Hsp70/40 machinery with a misfolded client is not sufficient to promote re-folding. The regulated cycle of client release and the potential for re-engagement is important, and this is dependent on nucleotide exchange. Indeed, changes in the availability or function of nucleotide exchange factors (NEFs) alone change client processing. Alterations in the NEF Sse1 were shown to alter yeast prion propagation in a strain-dependent manner^52^. Sse1 has been proposed to have multiple functions and can act as a disaggregase itself^53,54^. Our data suggest that the effect of the LGMDD1 mutants on the propagation of the [*RNQ+*] strain can be rescued by the deletion of Sse1 (HSP110). We also assessed the functional role of Sse1 in the rescue by using two Sse1 mutants; Sse1-K69Q and Sse1-G233D. We found a marked improvement in [*RNQ+*] prion propagation in yeast cells carrying these Sse1 mutants and harboring LGMDD1 mutants. Similarly, there was a considerable improvement in the refolding of luciferase activity in these cells. Notably, although the deletion of Sse1 alone showed partial improvement in [*RNQ+*] prion propagation in yeast cells, it did not show any difference in the refolding of luciferase, again perhaps indicating substrate specificity. Thus, this rescue further supports that the LGMDD1 mutants act in an Hsp70-dependent manner.

We hypothesize that the observed defects in the LGMDD1 mutants result in cellular phenotypes that are client-conformer specific. The Hsp70/DNAJ ATPase cycle is a process partitioned into two interconnected events; DNAJ is vital in the first half whereas NEFs play a significant role in the second half. Our data suggest that there are numerous defects associated with DNAJB6 (Sis1) mutants which result in either inhibition or delay in client processing in the first part of the Hsp70-ATPase cycle. Hsp70 has been shown to suppress proper substrate folding if it is not allowed to cycle off its client protein in various contexts^55,56^. Henceforth a longer interaction of LGMDD1 mutants with Hsp70 might lead to broader disruption of Hsp70-dependent processes, as this could titrate Hsp70 away from other clients^57^. Our data suggest that inhibiting the second half of the ATPase cycle, either by deletion or using Sse1 mutants, can have positive consequences on client processing. Interestingly, previous data suggest that the optimal NEF activity for protein disaggregation occurs at a reduced ratio of NEF:Hsp70 (1:10)^58–60^, and perhaps the deletion of Sse1 recapitulates such reduction in NEF activity in some manner (such as replacing the optimal NEF with another, such as Fes1). Moreover, the armadillo-type NEFs (budding yeast Fes1 and its human homolog HspBP1) employ flexible N-terminal release domains (RDs) with substrate-mimicking properties to ensure the efficient release of persistent substrates from Hsp70^61^. This is plausible due to the fact that NEFs perform dual functions: accelerating nucleotide exchange and securing Hsp70-liberated substrates. Of note, the high selectivity of exchange factors for their Hsp70 partner contributes to the functional heterogeneity of Hsp70 chaperone system^62^. These results indicate that fine-tuning of the two halves of the Hsp70 ATPase cycle involving LGMDD1 mutants and NEFs during the processing of its client proteins is critical (Fig. 8C). As such, we suggest that NEF inhibitors could provide a possible therapeutic strategy for the treatment of LGMDD1.

## Methodology

### Cloning, Expression and Purification of Recombinant Proteins

Sis1-WT, Sis1-mutants (F106L/N108L/D110Δ/F115I) and Ssa1-WT were cloned into pPROEx-Htb vector obtained from Addgene. The plasmid encodes a hexa-His-tag, a TEV cleavage site, and the respective cloned gene for expression. All Sis1 mutants were generated using the Quick Change Mutagenesis Kit (Agilent Technologies #200517). Primer sequences were generated using Agilent’s online primer design program. Mutagenesis was confirmed by sequencing the entire coding region of *SIS1.* Sis1-WT and Sis1-mutants were expressed at 16 °C, whereas Ssa1-WT was expressed at 18 °C to increase the fraction of soluble protein. All purification steps were carried out at 4 °C. Protein purity was more than 99% as determined by SDS/PAGE and Coomassie staining. Final protein concentration was estimated by Bradford assay, using bovine serum albumin as the standard. Following purification, all the proteins were frozen on liquid nitrogen and stored at −80 °C till further use. Sis1-WT and mutants were purified using standard protocol with some modifications. Briefly, these were purified from *Escherichia coli* strain Lemo 21(DE3) (New England Biolabs) grown in 2X YT medium at 30 °C until OD_600_ = 0.6–0.8. The cultures were induced with 0.5 mM IPTG and grown overnight at 16 °C. Cells were harvested and lysed in buffer A (50 mM Sodium phosphate buffer (pH 7.4), 300 mM NaCl, 5 mM MgCl_2_, 10 mM Immidazole, 0.1% Igepal, 0.01 M TCEP (tris(2-carboxyethyl)phosphine), protease inhibitor cocktail (EDTA-free from Roche) and a pinch of DNase I. Cell debris was cleared by centrifugation (20,000 *g*) and the supernatant loaded on cobalt-based Talon metal affinity resin. After washing, proteins were eluted as gradient fractions with buffer A containing increasing concentrations of imidazole (150 mM – 400 mM). Purified proteins were incubated with His-TEV (purified in the lab) protease at 30° for 1 h. The samples were extensively dialyzed at 4 °C and again passed through Talon metal affinity resin to remove the cleaved His tag and His-TEV protease. The pure proteins were concentrated and stored at −80 °C. Similarly, Ssa1-WT protein was also purified using an established procedure^63^. Briefly, protein was purified from *Escherichia coli* strain Rosetta 2(DE3) (Invitrogen) grown in LB medium with 300 mM NaCl at 30 °C until OD_600_ = 0.6–0.8. The culture was induced with 0.5 mM IPTG and grown overnight at 18 °C. Cells were harvested and lysed in buffer A (20 mM Hepes, 150 mM NaCl, 20 mM MgCl_2_, 20 mM KCl, protease inhibitor cocktail (EDTA-free from Roche)) using lysozyme. The rest of the protocol was similar to that of Sis1-WT and mutants, with only exception being that Ssa1-WT was eluted with buffer A containing 250 mM imidazole. Rnq1-WT full-length protein was purified exactly as described in previous publication from our lab^26^.

### Binding Assays

#### Substrate-binding ELISA assays

This was performed as described earlier with some modifications. Two different substrate proteins-Rnq1 and firefly luciferase (Promega Corporation) were denatured for 1 h at 25 °C in a buffer comprising of 3 M guanidine HCl, 25 mM HEPES (pH 7.5), 50 mM KCl, 5 mM MgCl_2_, and 5 mM DTT. Following denaturation, substrates were diluted in 0.1 M NaHCO_3_ and bound to microtiter plate (CoStar 3590 EIA plates, Corning) at a concentration of 0.4 μg/well for Rnq1 and 0.1 μg/well for luciferase, respectively. Unbound substrate was removed by washing with phosphate buffered saline (PBS). Unreacted sites were blocked with 0.2 M glycine (100 μl/well) for 30 min at 24 °C, followed by washing with PBS-T (PBS containing 0.05% Tween 20). Non-specific binding was eliminated by blocking with 0.5% fatty-acid-free bovine serum albumin (BSA) (Millipore Sigma) in PBS for 6 hours. The wells were subsequently washed with PBS-T. Sis1-WT and Sis1-mutants were serially diluted in PBST (substituted with 0.5% BSA) and incubated with substrate for overnight at 24 °C. After extensive washing with PBS-T, rabbit anti-Sis1 antibody (CosmoBio) at a dilution of (1:15000) was added and incubated for 2 h at 24 °C. This was followed by further washings and addition of donkey-anti-rabbit HRP-conjugated (Millipore Sigma) (1:4000) as secondary antibody. The amount of Sis1 retained was determined by developing a reaction using tetramethyl benzidine/H_2_O_2_ (TMB peroxidase EIA substrate) kit (Bio-Rad). The colour was measured at 450 nm (SpectraMax M2e fluorimeter microplate reader) after terminating the reaction with 0.02 N H_2_SO_4_.

##### Ssa1-binding assays

Ssa1 (200 nM) was immobilized in microtiter plate wells and dilutions of purified Sis1-WT and Sis1-mutants were incubated with it. Bound Sis1 was detected as described above.

#### Substrate bound Sis1 combined with Ssa1 binding assays

Denatured substrates (Rnq1 and luciferase) at a concentration of 0.4 ug/well and 0.1 ug/well, respectively, were mixed with Sis1-WT and Sis-mutant proteins (5 nM) and incubated for 1 h at 24 °C, prior to being adsorbed in the microtiter well plates. Following the steps of incubations, washings and blocking, serially diluted Ssa1-WT was added to each well. Subsequently, the wells were probed with rabbit-Ssa1 antibody (1:2000) (Abcam), followed by mouse anti-rabbit HRP conjugated secondary antibody (Millipore Sigma). The detection method used was similar to that described above for detecting Sis1.

#### Homo/Hetero dimeric nature of Sis1 determining binding assays

Serially diluted His-tagged cleaved Sis1-WT and Sis1-mutants were adsorbed in the microtiter well plates. Following washings, serially diluted uncleaved His-tagged Sis1-WT and Sis1-mutants were added to the wells in the following combinations [(Cleaved mutants + Uncleaved mutants =Homodimer); (Cleaved mutants + Uncleaved Sis1-WT = Heterodimer) along with appropriate controls for the assay. This was followed by washings, blocking and addition of mouse anti-His antibody (1:5000) (Invitrogen). Rabbit anti-mouse HRP conjugated antiserum (1:4000) (Millipore Sigma) was used as secondary antibody. The detection method was similar to that described above.

### Amyloid Fiber Formation and Thioflavin T kinetics

Purified Rnq1 was resuspended in 7 M guanidine hydrochloride and the protein concentration was determined. Rnq1 fibers were formed at 18°C, 25°C and 37°C with a starting monomer concentration of 8 μm in Fiber-formation buffer (FFB) (50 mM KPO_4_, 2 M Urea, 150 mM NaCl, pH 6). For the seeded kinetics experiments, the fibers were seeded using 5% (w/w) seed. The fiber formation and kinetics assays were performed in the presence of Thioflavin T dye and acid-washed glass beads (Sigma) for agitation as described earlier^26^. Kinetic assays of fiber formation were done in a SpectraMax M2e fluorimeter microplate reader. The change in Thioflavin-T fluorescence over time was measured using an excitation wavelength of 450 nm and emission wavelength of 481 nm.

### Colorimetric determination of ATPase activity

The ATPase assay was performed as described before^64^ with some modifications. Briefly, the ATPase reagent was made by combining 0.081% W/V Malachite Green with 2.3% W/V poly-vinyl alcohol, 5.7% W/V ammonium heptamolybdate in 6 M HCl, and water in 2:1:1:2 ratios (all purchased from Sigma with no further purification). This ATPase reagent was freshly prepared every day and was left standing for 2 h to get a stable green/golden solution, which was filtered through 0.45 μm syringe filters (Millipore Sigma) before use. ATPase activity in the absence of any client protein was tested by incubating Sis1-WT/mutants, Ssa1-WT in the ratio of (0.05:1.0 μM) with 1 mM ATP, in assay buffer (0.02% Triton X-100, 40 mM Tris–HCl, 175 mM NaCl, and 5 mM MgCl_2_, pH 7.5) at 37 °C for different time-intervals as indicated in the figure legends. At the end of incubation 25 μL of the reaction was added to a well in a 96 well plate, followed by 800uL of the ATPase reagent and 100 μL of 34% sodium citrate to halt any further ATP hydrolysis. The mixture was allowed to incubate for 30 min at 24 °C before absorbance at 620 nm was measured using a SpectraMax M2e fluorimeter microplate reader. A sample of ATP alone in buffer was treated exactly the same and was subtracted from the sample absorbance to account for intrinsic ATP hydrolysis. To account for variability in measurements a phosphate standard curve (using potassium phosphate) was created for each day of measurements. For ATPase activity in presence of client proteins, the same procedure was followed with only exception being the addition of client protein Rnq1 monomer (25 μM) and Rnq1 seeds (10% of which is used in final reaction) formed at three different temperatures (18°C, 25°C and 37°C) with the chaperones and ATP for incubation.

### Luciferase refolding assay

Heat denatured refolding of luciferase was performed as previously described^65^. Briefly, Ssa1 (2 μM) were incubated in refolding buffer (50mM Tris pH 7.4, 150 mM KCl, 5 mM MgCl_2_) supplemented with 1 mM ATP and an ATP regenerating system (10 mM phosphocreatine, 100 mg/ml phosphocreatine kinase) for 15 min at room temperature. Next, luciferase (25 nM) was added and incubated for a further 10 min. Then Sis1-WT/mutants (0.05 μM) was added and the mixture was heat shocked at 44 °C for 20 min. The reactions were then immediately moved to room temperature. Finally, 25 μL aliquots of the refolding reaction were then taken and added to 50 μL of luciferase assay reagent (Promega Corporation). At various time points, activity was then measured with a GloMax Luminometer (Promega Corporation).

### Titration assays

All the assays were performed by titrating the concentration of Sis1-mutants (F106L and F115I) with Sis1-WT such that the total concentration of protein was the same as been used individually across different assays.

### Yeast strains, plasmids and Transformation

The yeast strains used in this study are derived from *Saccharomyces cerevisiae* 74-D694 (*ade1-14 his3*-Δ200 *leu2-3, ll2 trpl-289 ura 3-52*). Yeast cells were grown and manipulated using standard techniques^66^. As indicated, cells were grown in rich media YPD (1% yeast extract, 2% peptone, 2% dextrose) or in synthetic defined (SD) media (0.67% yeast nitrogen base without amino acids, 2% dextrose) lacking specific nutrients to select for appropriate plasmids. Wild-type (WT) yeast harboring the s. d. medium [*RNQ*+] variant and the [*rnq-*] control strain were kindly provided by Dr. Susan Liebman (University of Nevada, Reno, Nevada, USA)^67^. Construction of *ΔSis1* [*rnq*-] and s. d. medium [*RNQ+*] yeast strains were described previously^12^. *ΔSse1* and all other delta strains in s. d. medium [*RNQ+*] background were created using the standard protocol. Medium containing 1mg/mL 5-fluoroorotic acid (5-FOA) that selects against cells maintaining *URA3-* marked plasmids was used to replace WT Sis1 with the mutant constructs using the plasmid shuffle technique. Plasmid transformations were done using polyethylene-glycol/lithium-acetate (PEG/LioAC) technique and the cells were selected using SD-trp/his plates.

Plasmid pRS316-*SIS1* was kindly provided by Dr. Elizabeth Craig (University of Wisconsin, Madison, Madison, Wisconsin, USA)^22^. Plasmids *pRS414-Sse1-K69Q,* and pRS414-*Sse1-G233D* were kind gifts from Dr. Kevin Morano (McGovern Medical School, UT Health, Houston, Texas, USA)^37^. Construction of pRS314-Sis1 and LGMDD1 mutants were described previously. Using the standard molecular techniques we constructed p413TEF-Sse1-K69Q and p413TEF-Sse1-G233D. Plasmid pRS316-GPD-Lux was a kind gift from Dr. Bernd Bukau (Center for Molecular Biology of Heidelberg University, Heidelberg, Germany)^39^.

### Protein fiber Transformation for Phenotypic analysis

Transformation of Rnq1 fibers into a [*rnq*-] 74-D694 (ade1-14, ura3-52, leu2-3,112, trp1-289, his3-200, sup35::RRP) yeast strain in the presence and absence of Sis1-WT/ mutants was conducted as described^68^. The resulting colonies formed after infecting fibers formed in vitro in the presence and absence of chaperones were replica plated onto rich medium (YPD) plates to assay for colony color. Colonies that appeared to have acquired the prion state by nonsense suppression were picked and spotted on YPD, YPD containing 3 mm GdnHCl, and SD-Ade for phenotypic analyses.

### Protein Analysis

Yeast samples were lysed with glass beads in buffer (100 mM Tris-HCl pH7.5, 200mM NaCl, 1mM EDTA, 5% glycerol, 0.5 mM DTT, 3 mM PMSF, 50 mM N-ethylmaleimide (NEM), complete protease inhibitor from Roche) and pre-cleared at 6000 rpm for 15 sec. Protein concentration of cells lysates was then normalized. For well-trap assays, samples were incubated at room temperature or 100°C in sample buffer (200mM Tris-HCl pH 6.8, 4% SDS, 0.4% bromophenol blue, 40% glycerol), then analyzed by SD-PAGE and western blot using an αRnq1 antibody. Boiled gel assays were performed as described previously^15^. Briefly, yeast cells were lysed with glass beads in buffer (25mM Tris-HCl pH7.5, 100mM NaCl, 1mM EDTA, protease inhibitors) and pre-cleared at 6,000 rpm for 1 minute at 4°C. Protein concentration of cell lysates was normalized using a Bradford assay and mixed with SDS-Page sample buffer (200mM Tris-HCl pH 6.8, 4% SDS, 0.4% bromophenol blue, 40% glycerol). Samples remained un-boiled and were loaded on a 12% polyacrylamide gel and run under constant current of 110V until the dye front migrated halfway through the resolving gel. The current was then stopped, and the gel in glass plates was sealed in plastic and boiled upright for 15 mins in a 95-100°C water bath. After boiling, gels were removed from the plastic cover and were reinserted in the SDS-PAGE apparatus, where voltage was re-applied until the dye migrated to the bottom of the gel. SDS-PAGE was followed by standard western blotting with αRnq1 antibody. Semi-denaturing agarose gel electrophoresis (SDD-AGE) for [*RNQ+*] fibers was performed as previously described^69^.

### Negative Staining Transmission Electron Microscopy

Rnq1 fibers were generated as described above. Negative staining of Rnq1 samples was performed by depositing 8uL of sample and incubating for one minute on carbon coated 200 mesh copper grids (01840-F, Ted Pella, Redding, CA), held by forceps carbon side up which had been freshly glow discharged for 30 seconds in a Solarus 950 plasma cleaner (Gatan, Pleasanton, CA). Post-incubation, each grid was washed five times in separate ultrapure water droplets and subsequently stained with 0.75% uranyl formate for 2 minutes. Excess uranyl formate was blotted off using filter paper (Whatman No.2, Fisher Scientific, Hampton, NH) and subsequently air dried. Sample grids were imaged using a JEOL JEM-1400 Plus Transmission Electron Microscope operating at an accelerating voltage of 120 kV equipped with an NanoSprint15 MKII sCMOS camera (AMT Imaging, Woburn, MA). Images were acquired using a total exposure time of 5 seconds containing ten 500ms drift frames which were subsequently aligned and averaged using the AMT ImageCapture Engine acquisition software at nominal magnifications ranging between 20,000-50,000x.

### Statistical analysis

Error bars represent standard error mean from at least three experiments. Significance was determined for two-sample comparisons using the unpaired *t*-test function with a threshold of two-tailed *p* values less than 0.05 for *, 0.01 for ** and 0.001 for ***.

## Acknowledgements

We are thankful to S. Liebman, J. Weissman, K. Morano, and E. Craig for plasmids and strains. We also thank Kevin Stein for initial experiments with Sse1. We are thankful to Patrick McConnell and Kevin M. Kaltenbronn for helping with protein purification and luciferase assay, respectively. We are thankful to members of True lab and Weihl lab for helpful discussions and comments on the manuscript. This work was supported by: National Institutes of Health Grant R01AR068797 to HLT and CCW. The funders had no role in study design, data collection and analysis, decision to publish, or preparation of the manuscript.

## Author contributions

A.K.B., C.C.W., and H.L.T. conceived and designed the experiments and analyzed data. A.K.B. performed all the experiments and wrote the initial draft. M.J.R. and J.A.J.F collected the negative-staining TEM images. J.A.D. performed the site-directed mutagenesis. A.K.B., C.C.W., and H.L.T. reviewed and edited the manuscript. All authors provided editorial input.

## References

1. Iyadurai, S. J. P. & Kissel, J. T. The Limb-Girdle Muscular Dystrophies and the Dystrophinopathies. Continuum (Minneap Minn) 22, 1954–1977 (2016).

2. Kley, R. A., Olivé, M. & Schröder, R. New aspects of myofibrillar myopathies. Curr Opin Neurol 29, 628–634 (2016).

3. Sarparanta, J. et al. Mutations affecting the cytoplasmic functions of the co-chaperone DNAJB6 cause limb-girdle muscular dystrophy. Nat Genet 44, 450–455, S1-2 (2012).

4. Harms, M. B. et al. Exome sequencing reveals DNAJB6 mutations in dominantly-inherited myopathy. Ann Neurol 71, 407–416 (2012).

5. Ruggieri, A. et al. Complete loss of the DNAJB6 G/F domain and novel missense mutations cause distal-onset DNAJB6 myopathy. Acta Neuropathol Commun 3, 44 (2015).

6. Palmio, J. et al. Novel mutations in DNAJB6 gene cause a very severe early-onset limb-girdle muscular dystrophy 1D disease. Neuromuscul Disord 25, 835–842 (2015).

7. Sato, T. et al. DNAJB6 myopathy in an Asian cohort and cytoplasmic/nuclear inclusions. Neuromuscul Disord 23, 269–276 (2013).

8. Hageman, J. et al. A DNAJB chaperone subfamily with HDAC-dependent activities suppresses toxic protein aggregation. Mol Cell 37, 355–369 (2010).

9. Watson, E. D., Geary-Joo, C., Hughes, M. & Cross, J. C. The Mrj co-chaperone mediates keratin turnover and prevents the formation of toxic inclusion bodies in trophoblast cells of the placenta. Development 134, 1809–1817 (2007).

10. Zhang, Y. et al. The Hsp40 family chaperone protein DnaJB6 enhances Schlafen1 nuclear localization which is critical for promotion of cell-cycle arrest in T-cells. Biochem J 413, 239–250 (2008).

11. Mitra, A., Menezes, M. E., Shevde, L. A. & Samant, R. S. DNAJB6 induces degradation of beta-catenin and causes partial reversal of mesenchymal phenotype. J Biol Chem 285, 24686–24694 (2010).

12. Stein, K. C., Bengoechea, R., Harms, M. B., Weihl, C. C. & True, H. L. Myopathy-causing mutations in an HSP40 chaperone disrupt processing of specific client conformers. J Biol Chem 289, 21120–21130 (2014).

13. Kampinga, H. H. & Craig, E. A. The HSP70 chaperone machinery: J proteins as drivers of functional specificity. Nat Rev Mol Cell Biol 11, 579–592 (2010).

14. Rose, J. M., Novoselov, S. S., Robinson, P. A. & Cheetham, M. E. Molecular chaperone-mediated rescue of mitophagy by a Parkin RING1 domain mutant. Hum Mol Genet 20, 16–27 (2011).

15. Pullen, M. Y., Weihl, C. C. & True, H. L. Client processing is altered by novel myopathy-causing mutations in the HSP40 J domain. PLoS One 15, e0234207 (2020).

16. Frost, B. & Diamond, M. I. Prion-like mechanisms in neurodegenerative diseases. Nat Rev Neurosci 11, 155–159 (2010).

17. Glover, J. R. & Lindquist, S. Hsp104, Hsp70, and Hsp40: a novel chaperone system that rescues previously aggregated proteins. Cell 94, 73–82 (1998).

18. True, H. L. The battle of the fold: chaperones take on prions. Trends Genet 22, 110–117 (2006).

19. Satpute-Krishnan, P., Langseth, S. X. & Serio, T. R. Hsp104-dependent remodeling of prion complexes mediates protein-only inheritance. PLoS Biol 5, e24 (2007).

20. Collinge, J. & Clarke, A. R. A general model of prion strains and their pathogenicity. Science 318, 930–936 (2007).

21. Palmio, J. et al. Mutations in the J domain of DNAJB6 cause dominant distal myopathy. Neuromuscul Disord 30, 38–46 (2020).

22. Lopez, N., Aron, R. & Craig, E. A. Specificity of class II Hsp40 Sis1 in maintenance of yeast prion [RNQ+]. Mol Biol Cell 14, 1172–1181 (2003).

23. Vitrenko, Y. A., Gracheva, E. O., Richmond, J. E. & Liebman, S. W. Visualization of aggregation of the Rnq1 prion domain and cross-seeding interactions with Sup35NM. J Biol Chem 282, 1779–1787 (2007).

24. Naiki, H. & Gejyo, F. Kinetic analysis of amyloid fibril formation. Methods Enzymol 309, 305–318 (1999).

25. Aron, R., Lopez, N., Walter, W., Craig, E. A. & Johnson, J. In vivo bipartite interaction between the Hsp40 Sis1 and Hsp70 in Saccharomyces cerevisiae. Genetics 169, 1873–1882 (2005).

26. Kalastavadi, T. & True, H. L. Analysis of the [RNQ+] prion reveals stability of amyloid fibers as the key determinant of yeast prion variant propagation. J Biol Chem 285, 20748–20755 (2010).

27. Yan, W. & Craig, E. A. The glycine-phenylalanine-rich region determines the specificity of the yeast Hsp40 Sis1. Mol Cell Biol 19, 7751–7758 (1999).

28. Perales-Calvo, J., Muga, A. & Moro, F. Role of DnaJ G/F-rich domain in conformational recognition and binding of protein substrates. J Biol Chem 285, 34231–34239 (2010).

29. Sondheimer, N., Lopez, N., Craig, E. A. & Lindquist, S. The role of Sis1 in the maintenance of the [RNQ+] prion. EMBO J 20, 2435–2442 (2001).

30. Wawrzynów, A. & Zylicz, M. Divergent effects of ATP on the binding of the DnaK and DnaJ chaperones to each other, or to their various native and denatured protein substrates. J Biol Chem 270, 19300–19306 (1995).

31. Bengoechea, R. et al. Inhibition of DNAJ-HSP70 interaction improves strength in muscular dystrophy. J. Clin. Invest. (2020) doi:10.1172/JCI136167.

32. Li, J., Qian, X. & Sha, B. Heat shock protein 40: structural studies and their functional implications. Protein Pept Lett 16, 606–612 (2009).

33. Sha, B., Lee, S. & Cyr, D. M. The crystal structure of the peptide-binding fragment from the yeast Hsp40 protein Sis1. Structure 8, 799–807 (2000).

34. Abrams, J. L., Verghese, J., Gibney, P. A. & Morano, K. A. Hierarchical functional specificity of cytosolic heat shock protein 70 (Hsp70) nucleotide exchange factors in yeast. J Biol Chem 289, 13155–13167 (2014).

35. Yakubu, U. M. & Morano, K. A. Roles of the nucleotide exchange factor and chaperone Hsp110 in cellular proteostasis and diseases of protein misfolding. Biol Chem 399, 1215–1221 (2018).

36. Shaner, L., Sousa, R. & Morano, K. A. Characterization of Hsp70 binding and nucleotide exchange by the yeast Hsp110 chaperone Sse1. Biochemistry 45, 15075–15084 (2006).

37. Shaner, L., Trott, A., Goeckeler, J. L., Brodsky, J. L. & Morano, K. A. The function of the yeast molecular chaperone Sse1 is mechanistically distinct from the closely related hsp70 family. J Biol Chem 279, 21992–22001 (2004).

38. Tipton, K. A., Verges, K. J. & Weissman, J. S. In vivo monitoring of the prion replication cycle reveals a critical role for Sis1 in delivering substrates to Hsp104. Mol Cell 32, 584–591 (2008).

39. Tessarz, P., Mogk, A. & Bukau, B. Substrate threading through the central pore of the Hsp104 chaperone as a common mechanism for protein disaggregation and prion propagation. Mol Microbiol 68, 87–97 (2008).

40. Smith, D. A., Carland, C. R., Guo, Y. & Bernstein, S. I. Getting folded: chaperone proteins in muscle development, maintenance and disease. Anat Rec (Hoboken) 297, 1637–1649 (2014).

41. Stein, K. C. & True, H. L. Extensive diversity of prion strains is defined by differential chaperone interactions and distinct amyloidogenic regions. PLoS Genet 10, e1004337 (2014).

42. Vaquer-Alicea, J. & Diamond, M. I. Propagation of Protein Aggregation in Neurodegenerative Diseases. Annu Rev Biochem 88, 785–810 (2019).

43. Caughey, B. & Kraus, A. Transmissibility versus Pathogenicity of Self-Propagating Protein Aggregates. Viruses 11, E1044 (2019).

44. Sanders, D. W. et al. Distinct tau prion strains propagate in cells and mice and define different tauopathies. Neuron 82, 1271–1288 (2014).

45. Mirbaha, H. et al. Inert and seed-competent tau monomers suggest structural origins of aggregation. Elife 7, e36584 (2018).

46. Sharma, A. M., Thomas, T. L., Woodard, D. R., Kashmer, O. M. & Diamond, M. I. Tau monomer encodes strains. Elife 7, e37813 (2018).

47. Deane, C. A. S. & Brown, I. R. Components of a mammalian protein disaggregation/refolding machine are targeted to nuclear speckles following thermal stress in differentiated human neuronal cells. Cell Stress Chaperones 22, 191–200 (2017).

48. Beissinger, M. & Buchner, J. How chaperones fold proteins. Biol Chem 379, 245–259 (1998).

49. Faust, O. et al. HSP40 proteins use class-specific regulation to drive HSP70 functional diversity. Nature 587, 489–494 (2020).

50. Rosenzweig, R., Nillegoda, N. B., Mayer, M. P. & Bukau, B. The Hsp70 chaperone network. Nat Rev Mol Cell Biol 20, 665–680 (2019).

51. Kampinga, H. H. et al. Function, evolution, and structure of J-domain proteins. Cell Stress Chaperones 24, 7–15 (2019).

52. Fan, Q., Park, K.-W., Du, Z., Morano, K. A. & Li, L. The role of Sse1 in the de novo formation and variant determination of the [PSI+] prion. Genetics 177, 1583–1593 (2007).

53. Shorter, J. The mammalian disaggregase machinery: Hsp110 synergizes with Hsp70 and Hsp40 to catalyze protein disaggregation and reactivation in a cell-free system. PLoS One 6, e26319 (2011).

54. Sadlish, H. et al. Hsp110 chaperones regulate prion formation and propagation in S. cerevisiae by two discrete activities. PLoS One 3, e1763 (2008).

55. Kirschke, E., Goswami, D., Southworth, D., Griffin, P. R. & Agard, D. A. Glucocorticoid receptor function regulated by coordinated action of the Hsp90 and Hsp70 chaperone cycles. Cell 157, 1685–1697 (2014).

56. Sekhar, A., Santiago, M., Lam, H. N., Lee, J. H. & Cavagnero, S. Transient interactions of a slow-folding protein with the Hsp70 chaperone machinery. Protein Sci 21, 1042–1055 (2012).

57. Park, S.-H. et al. PolyQ proteins interfere with nuclear degradation of cytosolic proteins by sequestering the Sis1p chaperone. Cell 154, 134–145 (2013).

58. Yakubu, U. M. & Morano, K. A. Suppression of aggregate and amyloid formation by a novel intrinsically disordered region in metazoan Hsp110 chaperones. J Biol Chem 296, 100567 (2021).

59. Dragovic, Z., Broadley, S. A., Shomura, Y., Bracher, A. & Hartl, F. U. Molecular chaperones of the Hsp110 family act as nucleotide exchange factors of Hsp70s. EMBO J 25, 2519–2528 (2006).

60. Wentink, A. S. et al. Molecular dissection of amyloid disaggregation by human HSP70. Nature 587, 483–488 (2020).

61. Gowda, N. K. C. et al. Nucleotide exchange factors Fes1 and HspBP1 mimic substrate to release misfolded proteins from Hsp70. Nat Struct Mol Biol 25, 83–89 (2018).

62. Brehmer, D. et al. Tuning of chaperone activity of Hsp70 proteins by modulation of nucleotide exchange. Nat Struct Biol 8, 427–432 (2001).

63. Sharma, D. & Masison, D. C. Single methyl group determines prion propagation and protein degradation activities of yeast heat shock protein (Hsp)-70 chaperones Ssa1p and Ssa2p. Proc Natl Acad Sci U S A 108, 13665–13670 (2011).

64. Chang, L. et al. High-throughput screen for small molecules that modulate the ATPase activity of the molecular chaperone DnaK. Anal Biochem 372, 167–176 (2008).

65. Lindstedt, P. R. et al. Enhancement of the Anti-Aggregation Activity of a Molecular Chaperone Using a Rationally Designed Post-Translational Modification. ACS Cent Sci 5, 1417–1424 (2019).

66. Gietz, D., St Jean, A., Woods, R. A. & Schiestl, R. H. Improved method for high efficiency transformation of intact yeast cells. Nucleic Acids Res 20, 1425 (1992).

67. Bradley, M. E., Edskes, H. K., Hong, J. Y., Wickner, R. B. & Liebman, S. W. Interactions among prions and prion ‘strains’ in yeast. Proc Natl Acad Sci U S A 99 Suppl 4, 16392–16399 (2002).

68. Tanaka, M. & Weissman, J. S. An efficient protein transformation protocol for introducing prions into yeast. Methods Enzymol 412, 185–200 (2006).

69. Bardill, J. P. & True, H. L. Heterologous prion interactions are altered by mutations in the prion protein Rnq1p. J Mol Biol 388, 583–596 (2009).

70. Vitrenko, Y. A., Pavon, M. E., Stone, S. I. & Liebman, S. W. Propagation of the [PIN+] prion by fragments of Rnq1 fused to GFP. Curr Genet 51, 309–319 (2007).

