## Supplemental figures for "Myopathy mutations in DNAJB6 slow conformer specific substrate processing that is corrected by NEF modulation": Supplementary Figures.pdf

**Supp. Fig. 1**

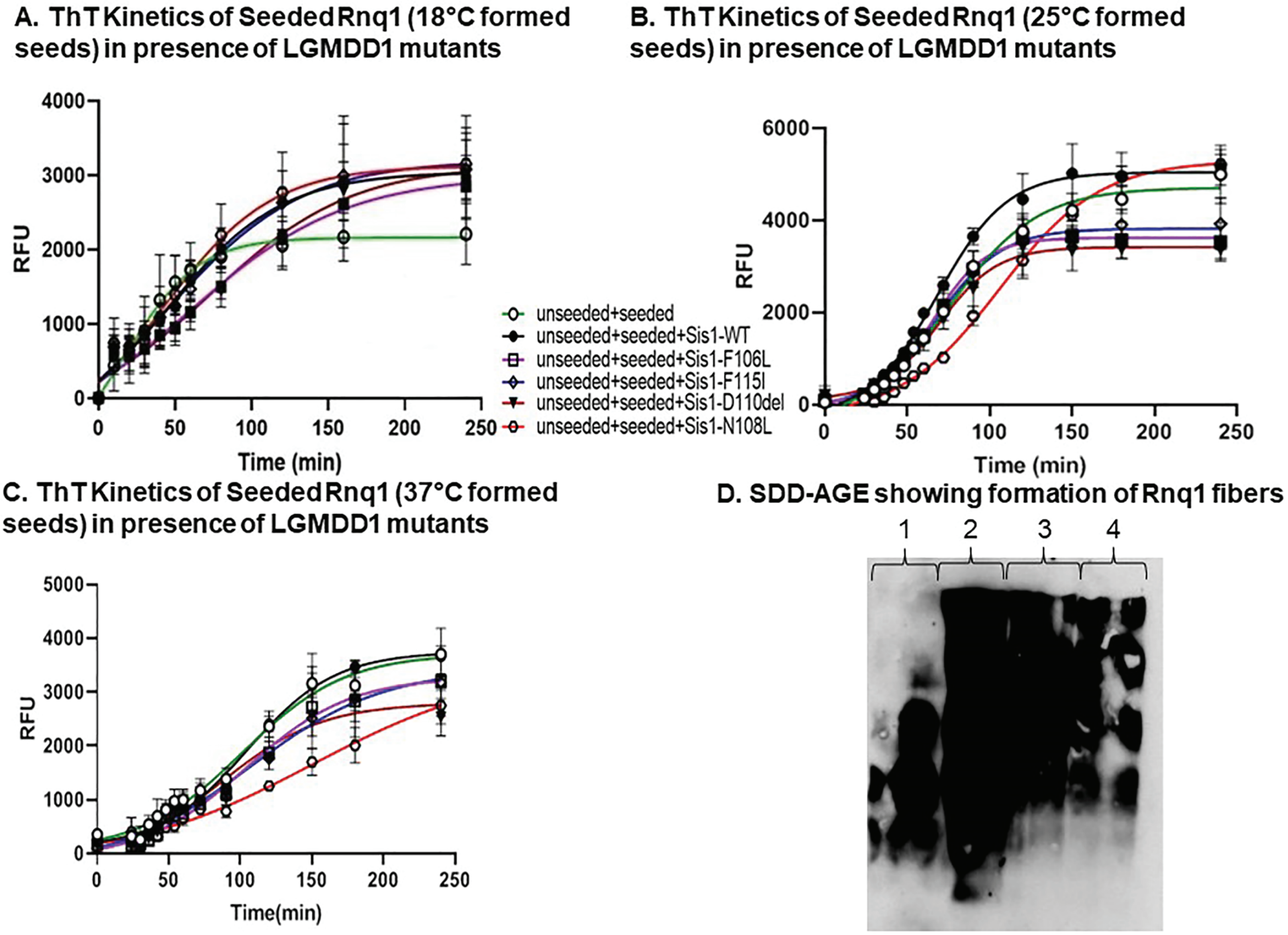


Supplementary Figure 1. **LGMDD1 G/F domain mutants behave differentially in aggregation assays with preformed Rnq1 seeds.** (A), (B), and (C) Kinetics of Rnq1 fibrillation in the presence of unseeded and seeded Rnq1 only **(green)**, Sis1-WT **(black)**, Sis1-F106L (**purple**), Sis1-N108L (**red**), Sis1-D110Δ (**brown**), or Sis1-F115I (**blue**) was measured by Th-T fluorescence assay. Note the abrupt increase in the fluorescence intensity of Th T indicates the formation of Rnq1 aggregates. Additionally, the fluorescence intensity of the dye achieved a plateau, indicating the stationary phase in fibrillation of Rnq1. *Inset*:-Fitted graph using the aggregation kinetics equation $y\ \ y\ i\ m\ x + (y\ f\ m\ x\ f\ \ \ e^{\ x\ \ \tau})$ where (yi + mxi) is the initial line, (yf + mxf) is the final line and x0 is the midpoint of maximum signal". 10 % of seeds (v/v) formed at different temperatures 18°C, 25°C, and 37°C were used in the assay. (D) SDD-AGE showing the formation of Rnq1 fibers at different temperatures. Lane 1: Rnq1 monomer, Lane 2-4: Rnq1 fibers formed at 18°C, 25°C, and 37°C respectively. Higher molecular weight aggregates formed by Rnq1 fibers appear as a smear in lanes 2, 3, and 4.

### Supp. Fig. 2

#### A. Cartoon diagram showing fiber transformation assay for selecting [*RNQ*+] variant distribution

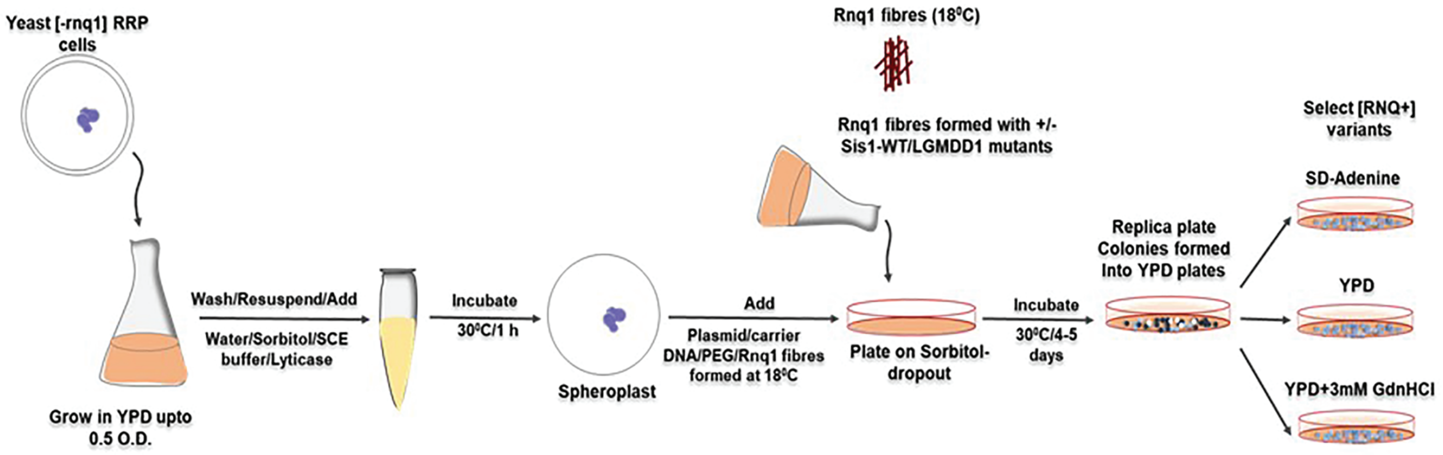


Supplementary Figure 2. **Cartoon depiction of the protein fiber transformation assay and the prion strain scoring method.** We took advantage of our [*RNQ*+] reporter protein (RRP) to phenotypically assess [*RNQ*+] prion variants[69]. RRP is a chimeric protein with the PFD of Rnq1 (amino acids 153–405) fused to the translation termination domain of Sup35 (amino acids 124–685)[70]. RRP co-aggregates with Rnq1 in the [*RNQ*+] state, resulting in prion-dependent nonsense suppression and thereby providing a phenotypic assay for the [*RNQ*] state. The level of nonsense suppression afforded by RRP changes with [*RNQ*+] prion variants, enabling us to distinguish variants by phenotype. Our yeast strain harbors an adenine mutation (*ade1-14*) and this leads to red coloration of colonies grown in rich medium (YPD) that are efficient in translation termination. The aggregation of RRP in different [*RNQ*+] variants result in different levels of suppression of the *ade1-14* premature stop codon and corresponding changes in the production of Ade1.

Supp. Fig. 3

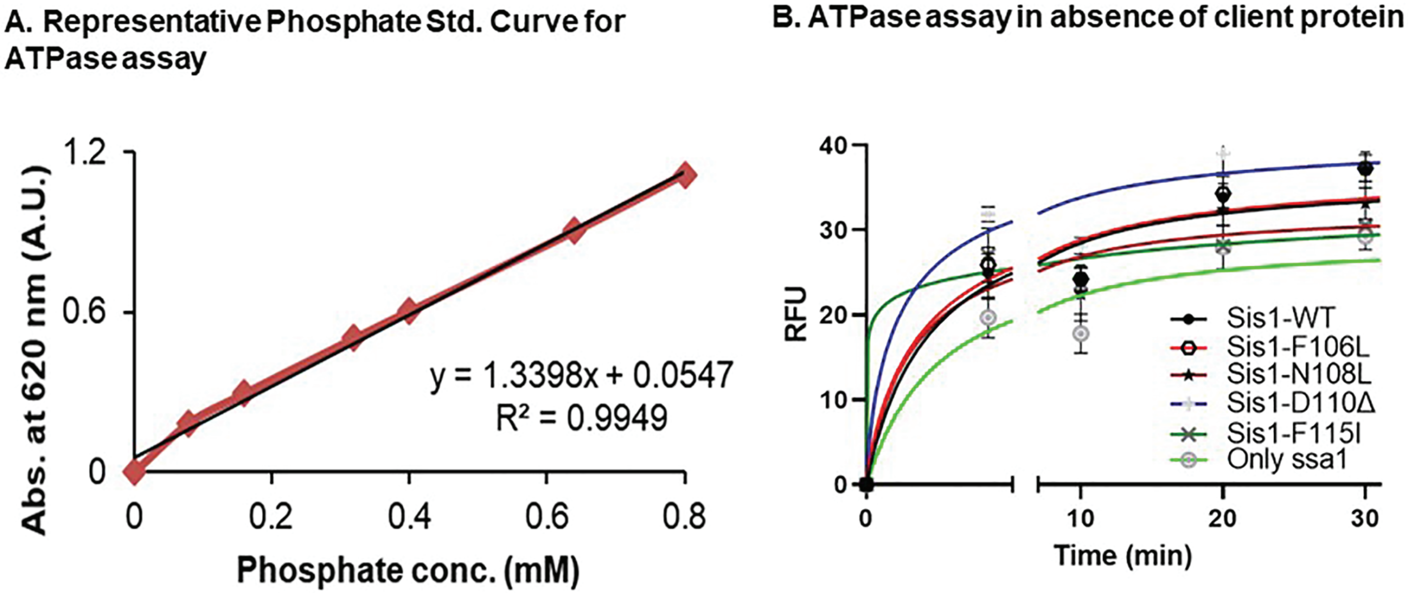


Supplementary Figure 3. (A) Representative Phosphate Standard Curve. (B) ATPase assay in absence of client protein. Sis1-WT or mutants (0.05 μM) and Ssa1 (1 μM) combined with ATP (1 mM).

**Supp. Fig. 4**

**Binding efficiency of LGMDD1 mutants with Ssa1-WT in absence of client substrate**

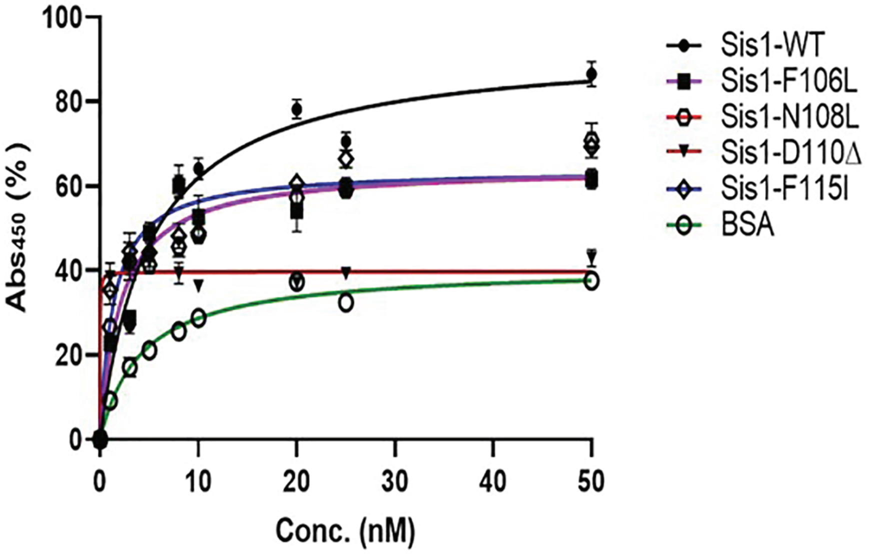


Supplementary Figure 4. **Association between LGMDD1 G/F domain mutants and Ssa1-WT was altered and is independent of client interaction**. Ssa1 (200 nM) was immobilized in microtiter plate wells and dilutions (0, 1, 3, 5, 8, 10, 20, 25, 50 nM) of purified Sis1-WT (**black**), and Sis1-mutants (Sis1-F106L (**purple**), Sis1-N108L (**red**), Sis1-D110Δ (**brown**), or Sis1-F115I (**blue**)) or BSA **(green)** as a control, were incubated with it. Bound Sis1 was detected using an αSis1 antibody.

**Supp. Fig. 5**

**A. ATPase assay by titrating F106L concentration**

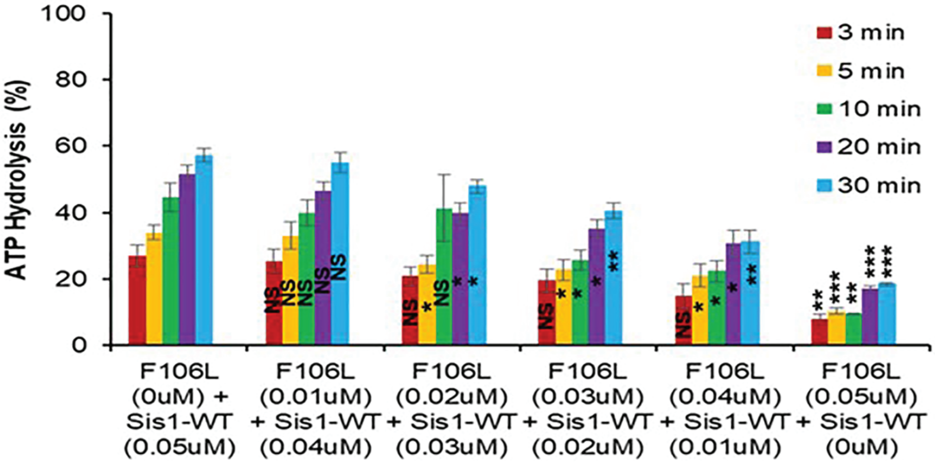


**B. ATPase assay by titrating F115I concentration**

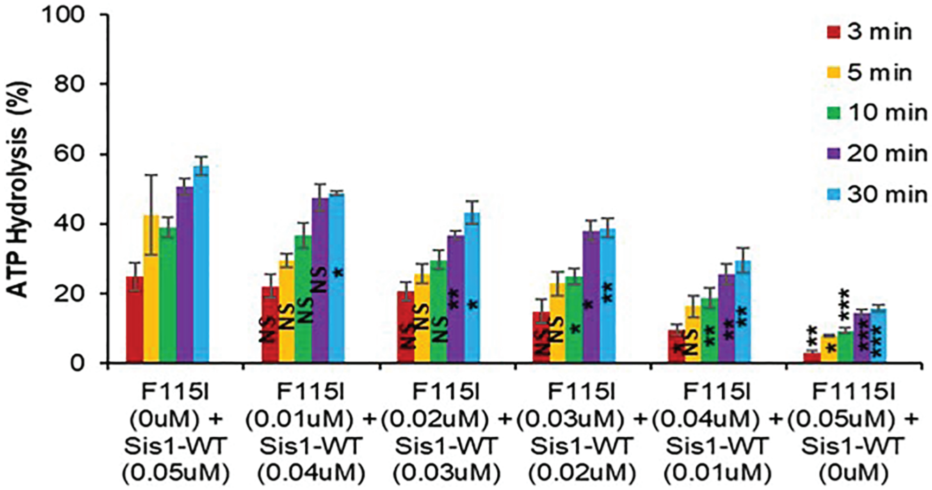


Supplementary Figure 5. **LGMDD1 G/F domain mutants inhibit Sis1-WT induced ATPase activity of Ssa1.** Stimulation of Ssa1 ATPase activity in the presence of Rnq1 seeds formed at 25°C. A total of Ssa1 (1 µM) in complex with ATP (1 mM) in the presence of Sis1-WT or mutants Sis1-mutants (Sis1-F106L (**A**), or Sis1-F115I (**B**)) (0.05 µM) or in the combination of half and half with Sis1-WT and mutants (0.5 µM each) were added. Fraction of ATP converted to ADP was determined at 3 min (**brown**), 5 min (**yellow**), 10 min (**green**), 20 min (**purple**), and 30 min (**blue**). A total of 10% seeds (v/v) were used in each reaction.

**Supp. Fig. 6**

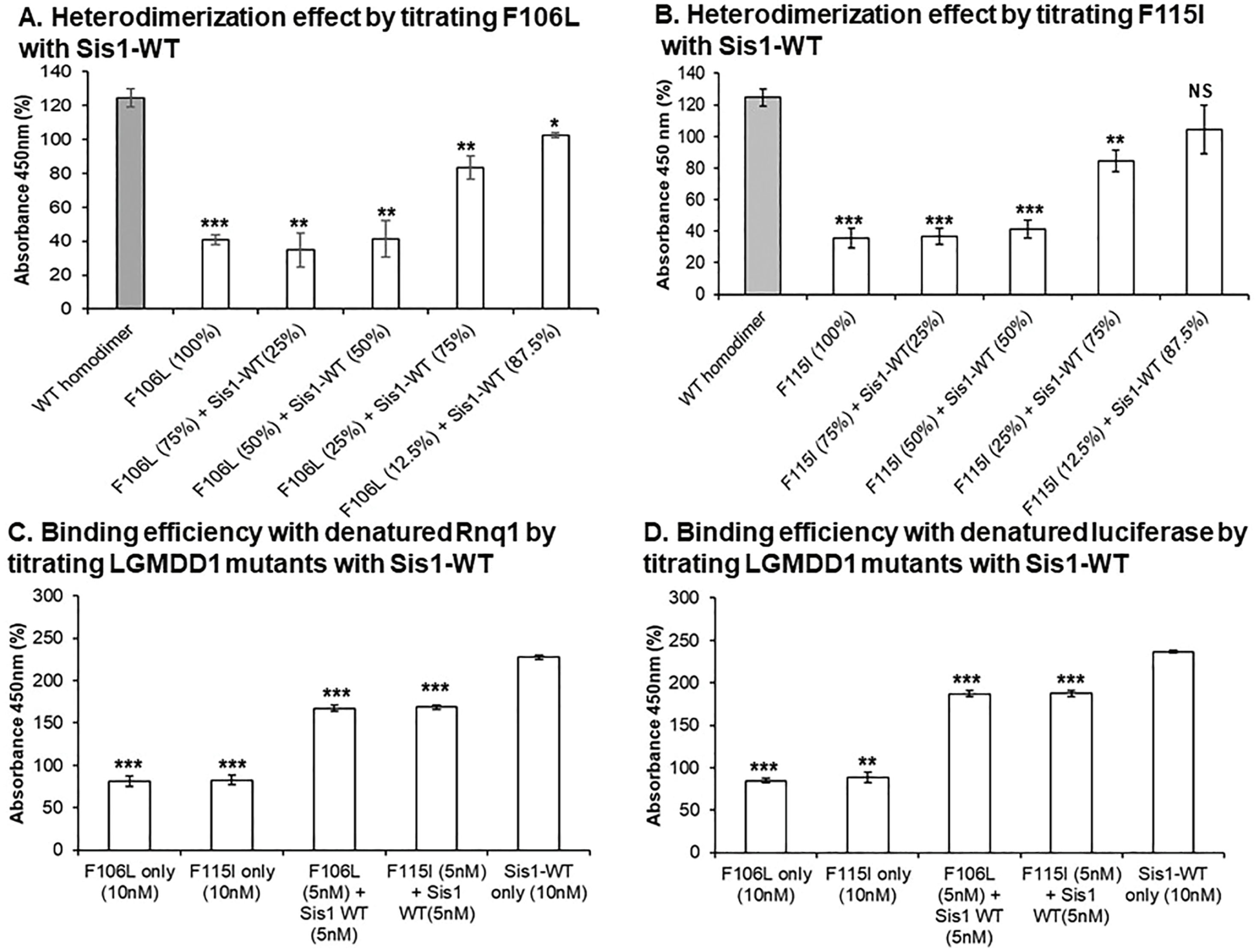


Supplementary Figure 6. **LGMDD1 G/F domain mutants inhibit the ability of wild-type Sis1 to dimerize and bind substrates.** Sis1-WT was used in one row of the microtiter plate (as homodimer control, **black**) and the rest rows were titrated with different concentrations of (A) F106L, and (B) F115I. Values were compared with Sis1-WT homodimer (**grey bar**). Binding of purified Sis1-WT (10 mM), Sis1-F106L (10 mM), Sis1-F115I (10 mM), or equimolar Sis1-WT and mutants (5 mM each) to denatured Rnq1 (C) and luciferase (D). Here, values were compared with Sis1-WT only.

Supp. Fig. 7

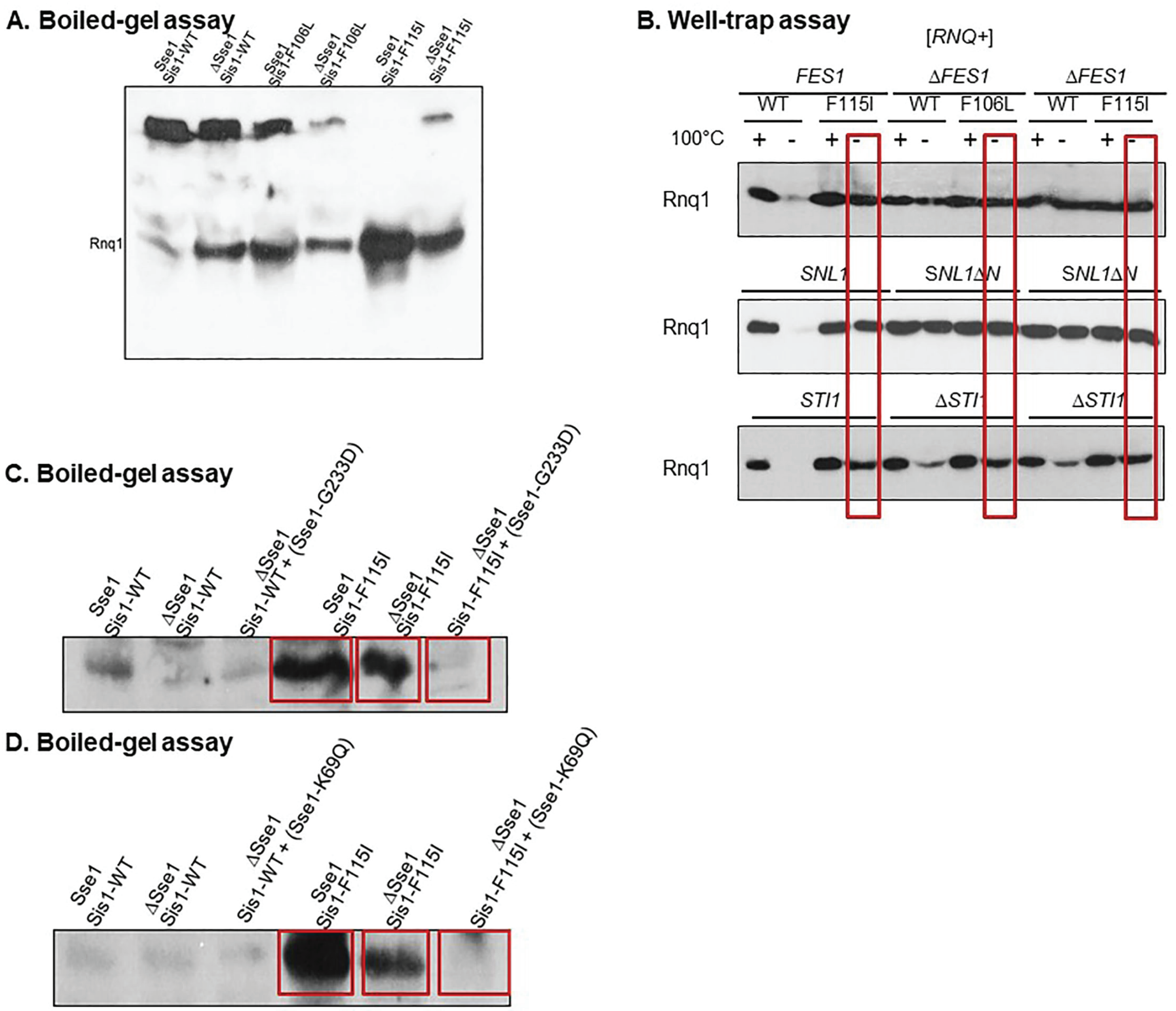


Supplementary Figure 7. **Deletion Sse1 function partially rescues prion propagation of [*RNQ*+].** (A) Boiled gel assays show the presence of more aggregated and less soluble Rnq1 protein in Sse1 cells expressing Sis1-WT, indicating proper [RNQ+] propagation. Notice that the prion propagation was altered in Sse1 cells with the expression of LGMDD1 mutants, however, were partially restored with the loss of Sse1 in these cells. (B) Deletion of the other NEF's did not rescue the propagation of [*RNQ*+]. (C) and (D) Sse1 mutants rescue prion propagation of [*RNQ*+]. Boiled gel assays showed less soluble Rnq1 protein (just like in Sse1 cells harboring Sis1-WT) in Δ*Sse1* cells co-expressing LGMDD1 mutants and (Sse1-G233D) and (Sse1-K69Q) respectively, indicating restoration of [*RNQ*+] prion propagation.
